# Nuclear assembly in giant unilamellar vesicles encapsulating *Xenopus* egg extract

**DOI:** 10.1101/2024.06.25.600006

**Authors:** Sho Takamori, Hisatoshi Mimura, Toshihisa Osaki, Tomo Kondo, Miyuki Shintomi, Keishi Shintomi, Miho Ohsugi, Shoji Takeuchi

## Abstract

The reconstitution of a cell nucleus in a lipid bilayer-enclosed synthetic cell makes great strides in bottom-up synthetic biology. In this study, we propose a method for assembling a nucleus in giant unilamellar vesicles (GUVs). To induce reconstitution of the nucleus, we utilize interphase egg extract of African clawed frogs *Xenopus laevis*, known as a biochemically controllable cell-free system capable of transforming an added sperm chromatin into a nucleus *in vitro*. We enhanced GUV formation efficiency by the inverted emulsion method through incorporating prolonged waiting time and adding chloroform into lipid-dispersed oil, facilitating subsequent nuclear assembly reactions in the GUVs.Characterization of nucleus-like structures formed in the GUVs revealed the presence of dense DNA and accumulated GFP-NLS in the structure, indicative of functional nuclear import. Immunostaining further validated the presence of nuclear pore complexes on the surfaces of these nucleus-like structures. Our approach offers a versatile platform for constructing artificial cellular systems that closely mimic eukaryotic cells.

**Teaser:** A cell nucleus is reconstituted in lipid bilayer-enclosed confinements using egg extract from African clawed frogs.

## Introduction

Cell nucleus serves as a crucial subcellular compartment in eukaryotic cells, responsible for overseeing the maintenance of genetic materials and orchestrating their functions. Recent advancements in bottom-up synthetic biology have focused on constructing or integrating cell-like functional subcellular compartments in synthetic cells (*1-5*). These efforts aim to replicate the functions of natural subcellular compartments such as photosynthesis in chloroplasts (*6-8*), mitochondrial respiration (*9-11*), and spatial organization of gene expression reactions in the nucleus (*12-13*). Among various eukaryotic subcellular compartments, the nucleus stands out as a central element in the regulation of eukaryotic cellular processes. Constructing the nucleus in synthetic cells encapsulated with a lipid bilayer represents a significant stride towards developing artificial eukaryotic cellular structures (*14*). Reconstitution of natural cell nucleus has previously been achieved in bulk solution (*15-17*), microfluidic devices (*18,19*), and water-in-oil emulsions (*20-22*), by utilizing egg extracts of the African clawed frog *Xenopus laevis*. The “interphase” egg extract of *Xenopus laevis* is known as a biochemically manipulable cell-free system, capable of transforming sperm chromatin into a nucleus *in vitro* (*15-17*). However, the challenge of assembling the nucleus in a lipid bilayer-enclosed confinement remains unmet. While encapsulation of an unspecified type of the egg extract into lipid bilayer-enclosed confinement has previously been reported (*23*), the typical yield in encapsulation of the interphase egg extract is low. This low yield can be attributed to the reported destabilization of the lipid bilayer in the presence of *Xenopus* egg extract (*24*), highlighting the challenge of achieving stability in the lipid membrane surrounding the egg extract. Thus, stabilizing the lipid bilayer could significantly improve the efficiency of encapsulation and enable the reconstitution of a nucleus in cell-like confinement enclosed with a lipid membrane.

In this study, we reconstitute a cell nucleus in giant unilamellar vesicles (GUVs). To accomplish the assembly of a nucleus in GUVs, we confine interphase egg extract of *Xenopus laevis* supplemented with *Xenopus* sperm chromatin by the inverted emulsion method. We incorporate an extended waiting time as a parameter to enhance GUV yield, based on the finding that the formation of a lipid monolayer at the oil-water interface, a crucial step in the inverted emulsion method, often requires a prolonged period (*25*). Additionally, we introduce chloroform into our lipid-dispersed oil, following reported procedures that involve adding organic solvents to promote GUV formation by aggregating lipids and improving their adsorption rate at the oil-water interface (*26*). Using GUVs formed by encapsulating the interphase egg extract, we induce nuclear assembly reactions in the GUVs. To confirm nuclear reconstitution, we first identify the functionality of the nuclear membrane by a nuclear import assay. Then, we conduct molecular confirmation of the nuclear membrane by immunostaining, and quantitatively characterize the nuclear assembly events and geometric features of the formed GUVs and nuclei.

## Results

### Formation of GUVs encapsulating *Xenopus* egg extract

To improve the encapsulation of *Xenopus* interphase egg extract (see “Materials and Methods” for details on the preparation of *Xenopus* interphase egg extract and sperm chromatin) into GUVs by the inverted emulsion method, we assessed the impact of implemented waiting time (*τ*: duration of waiting time; *τ* = 0/60/120 min) and added chloroform (*Δ*: relative volume of added chloroform to the lipid-dispersed oil; *Δ* = 0/5/10%) in our protocol of the inverted emulsion method (Fig. 1; see “Materials and Methods” for details on the intra-GUV nuclear assembly protocol). Representative confocal micrographs of the formed GUVs are shown in Fig. 2A. At *τ* = 0 min and *Δ* = 0%, we observed that the number and size of GUVs encapsulating the egg extract were notably small, unlike the GUVs encapsulating a typical dilute aqueous solution shown in Fig. S1. Meanwhile, we found an increasing trend in the number of GUVs encapsulating the egg extract with prolonged waiting time and higher volumes of added chloroform. These observations suggest that GUV-formation via the inverted emulsion method can be enhanced by implementing prolonged waiting time and adding chloroform to lipid-dispersed oil.

**Fig. 1.**
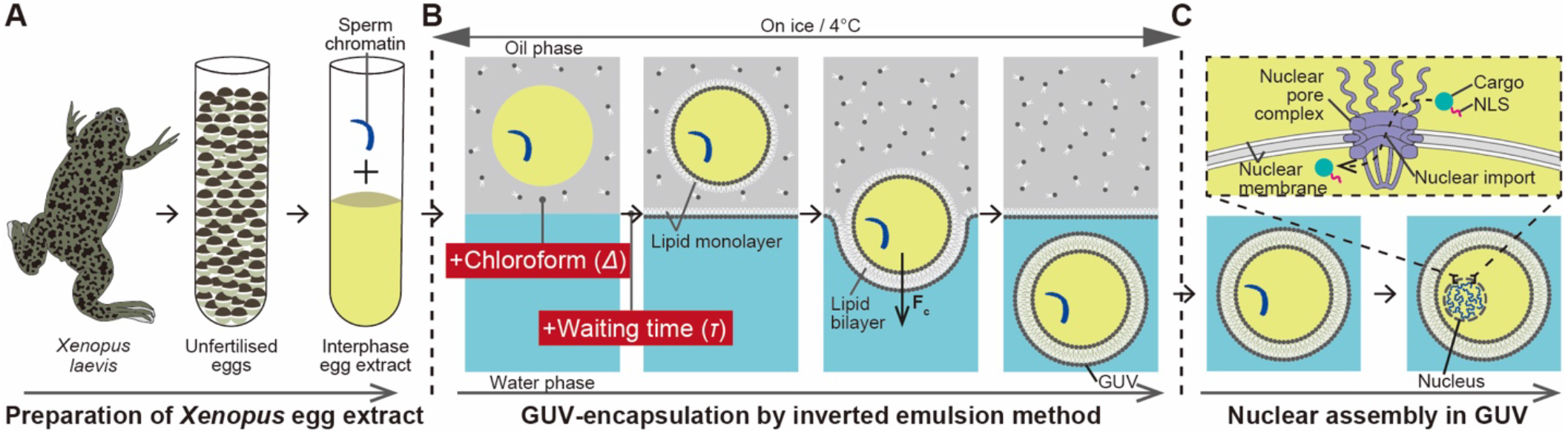
Three-step protocol for the assembly of a cell nucleus in GUVs. (**A**) Preparation of *Xenopus* interphase egg extract. Sperm chromatin is added to the egg extract prior to encapsulation into GUVs. (**B**) Encapsulation of sperm chromatin-added interphase egg extract into GUVs by the inverted emulsion method. Lipid monolayers are formed at the interfaces between water and oil phases. The formation of a lipid monolayer at the interfaces is enhanced by incubating the interfaces for the duration of the waiting time (*τ*) and adding chloroform in a volume (*Δ*) into lipid-dispersed oil. Centrifugal force (F_c_) allows the egg extract emulsions surrounded by a lipid monolayer to pass through a lipid monolayer formed at the interface between the oil phase and the outside water phase, transforming them into GUVs. (**C**) Induction of nuclear assembly reactions in GUVs by incubation at 22/23°C for 90 min.

**Fig. 2.**
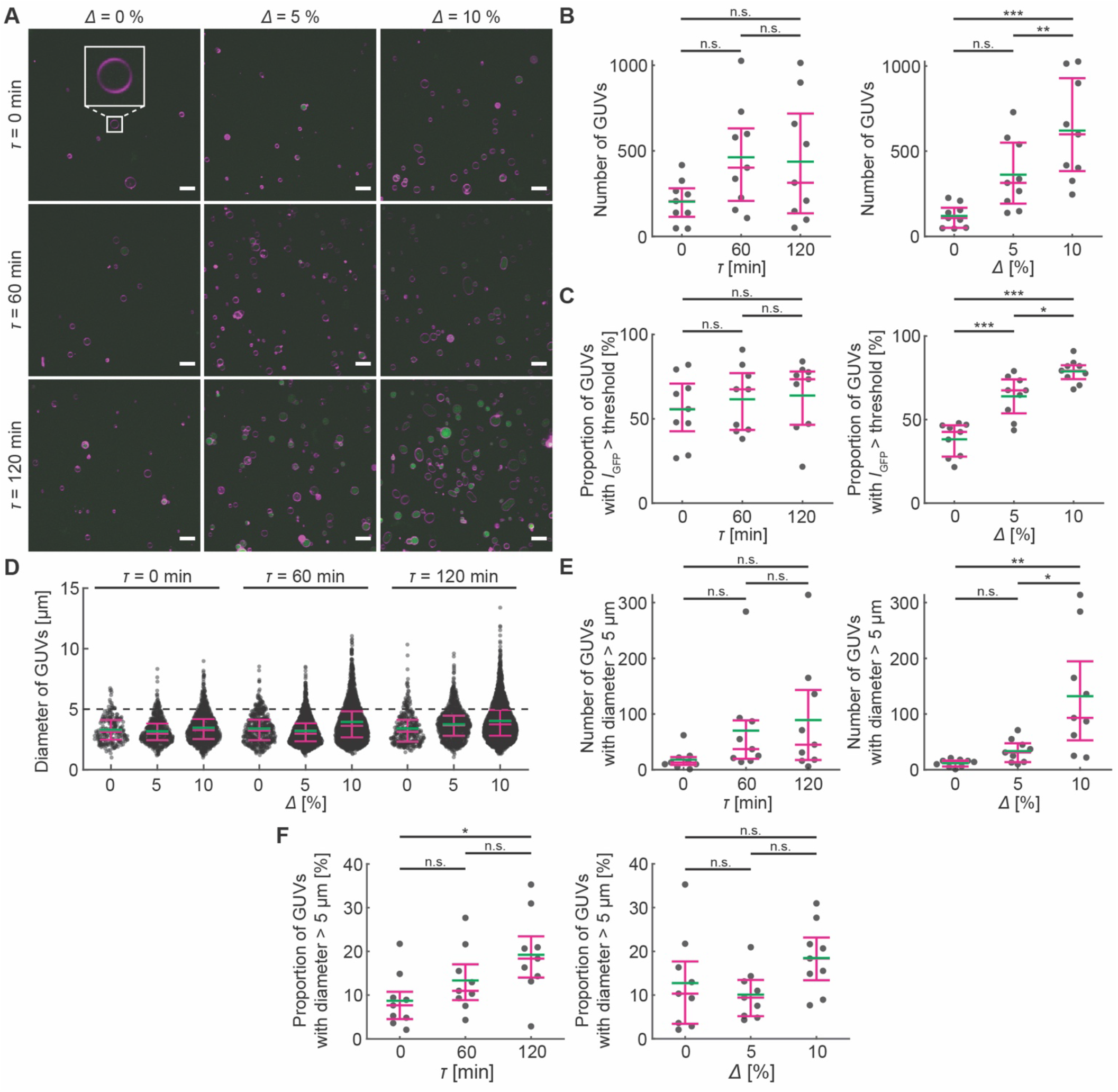
Formation of GUVs encapsulating *Xenopus* interphase egg extract. (**A**) Representative micrographs of GUVs encapsulating the sperm chromatin-added interphase egg extract, formed using the inverted emulsion method. The symbol *τ* denotes the duration of waiting time, and *Δ* denotes the volume of chloroform added to the lipid-dispersed oil. Magenta indicates lipid membrane (RhPE), and green indicates GFP dispersed in the egg extract. An inset shows a zoom-in of a typical GUV formed at *τ* = 0 min and *Δ* = 0%. Scale bar: 10 μm. (**B**) Number of GUVs plotted against *τ* (left) and *Δ* (right). (**C**) Proportion of GUVs with mean GFP intensity exceeding the computed threshold among the observed GUVs. (**D**) Diameter of all GUVs. (**E**) Number of GUVs with diameter larger than 5 μm. (**F**) Proportion of GUVs with diameter larger than 5 μm among the observed GUVs. Bars represent the mean (green) and Q1-Q3 quartiles (magenta). See “Materials and Methods” for the details on statistical analysis and Tables S1-12 for the results.

To evaluate the impact of waiting time and chloroform addition, we analyzed 1) the number of GUVs, 2) encapsulation efficiency of *Xenopus* egg extract in the GUVs, and 3) GUV diameters. We found that both waiting time and chloroform volume significantly increased the number of GUVs formed, with no significant interactions between these parameters (Table S1). Pairwise comparisons highlighted the role of chloroform volume, especially between *Δ* = 0% vs. 10% and *Δ* = 5% vs. 10%, while waiting time had no clear contribution (Tables S2-3, Fig. 2B). Chloroform volume notably influenced the proportion of GUVs encapsulating GFP-dispersed egg extract, enhancing encapsulation across all *Δ* conditions, whereas waiting time had no significant effect (Tables S4-6, Fig. 2C). Although most GUVs were below 5 μm (Fig. 2D), both waiting time and chloroform volume significantly increased the number of GUVs larger than 5 μm, with chloroform having a more distinct effect (Tables S7-9, Fig. 2E). For the percentage of GUVs with diameters larger than 5 μm, only waiting time had a significant effect (Table S10), increasing the proportion between *τ* = 0 min and 120 min, while chloroform volume had no noticeable impact (Tables S11-12, Fig. 2F). Overall, these results suggest that *i*) waiting time and chloroform volume both increase the number of GUVs and the number of larger GUVs, *ii*) chloroform enhances encapsulation of the egg extract, and *iii*) waiting time increases the proportion of larger GUVs.

### Formation of nucleus-like structures in GUVs

To initiate nuclear assembly reactions in GUVs encapsulating sperm chromatin-added *Xenopus* interphase egg extract, GUVs formed under various waiting time and chloroform conditions were incubated at 22ºC (or 23ºC) for 90 min. A nuclear import assay utilizing GFP fused with a nuclear localization signal derived from simian vacuolating virus 40 (GFP-NLS; see “Materials and Methods” for details on the preparation of GFP-NLS) was conducted as the primary means of identifying nucleus assembly, following established protocols in water-in-oil emulsions (*20-22*). In this assay, GFP-NLS is expected to be transported into the nucleus if a functional nuclear membrane with nuclear pore complexes is assembled on the surface as illustrated in Fig. 1C, while passive dispersion of GFP without the NLS-peptide is anticipated in the GUV lumen. Hereafter, the former condition is denoted as “NLS(+)”, and the latter as “NLS(-)”. Representative micrographs of these GUVs in the top row of Fig. 3A and Fig. S2 demonstrated that GFP clearly accumulated at the same location as dense DNA in the GUVs under the NLS(+) condition, suggesting active nuclear import of GFP-NLS into the assembled nucleus. In contrast, GFP was homogeneously dispersed in the GUVs containing dense DNA under the NLS(-) condition, albeit with a slight accumulation of GFP lacking the NLS-peptide at the same location as dense DNA observed in several confocal micrographs of GUVs in NLS(-) samples (Fig. S3). These confocal micrographs indicate successful assembly of nucleus-like structures in GUVs.

**Fig. 3.**
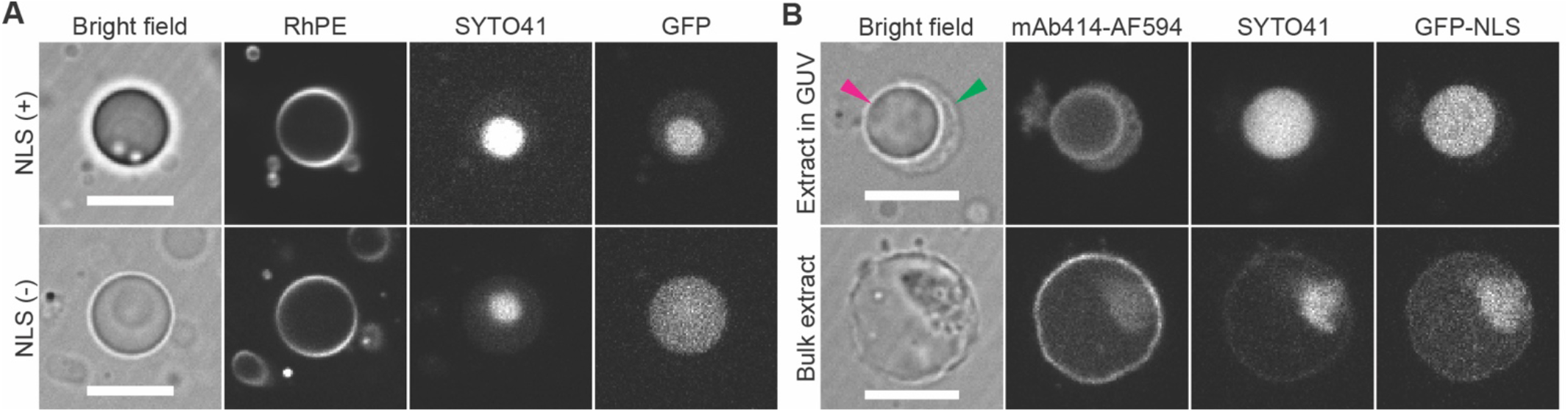
Formation of nucleus-like structures in GUVs and immunostaining of nuclear pore complexes. (**A**) GUVs after incubation for nuclear assembly reactions. RhPE: Lipid membrane, SYTO 41: nucleic acids (mainly DNA) stain. NLS(+): GFP with attached nuclear localization signals (NLS); NLS(-): GFP without NLS peptide. See Figs. S2-3 for additional examples. (**B**) A GUV containing a nucleus-like structure fixed with 2% glutaraldehyde and immunostained with mAb414-AF594 (top), and a nucleus formed under bulk condition, fixed and immunostained similarly (bottom). Arrowheads indicate GUV membrane (green) and nucleus-like structure (magenta). See Figs. S4-5 for additional examples. Scale bar: 10 μm.

### Immunostaining of nuclear pore complexes on nuclear membrane

To verify the presence of a nuclear membrane on the surface of nucleus-like structures formed in GUVs (top row of Fig. 3A and Fig. S2), we conducted immunostaining of nuclear pore complexes (NPCs) using NPC antibodies (mAb414) (*27,28*). GUVs containing a nucleus-like structure were fixed with 2.0% glutaraldehyde and stained with mAb414-Alexa Fluor 594 (mAb414-AF594; see “Materials and Methods” for the details on immunostaining). Confocal micrographs of the GUVs are shown in the top row of Fig. 3B and Fig. S4. As a positive control experiment, we conducted immunostaining of nuclei formed in the bulk form of sperm chromatin-added *Xenopus* interphase egg extract using the same procedure, as depicted in the bottom row of Fig. 3B and Fig. S5. These confocal micrographs confirmed accumulations of mAb414-AF594 signals at the periphery of nucleus-like structures in the GUVs, as well as at the periphery of nuclei formed in the bulk extract. Manual inspection of the micrographs revealed that more than 15% of nucleus-like structures showed the presence of NPCs on their surfaces (28 nucleus-like structures among the 169 structures fixed, immunostained, and inspected in total). The rest of nucleus-like structures exhibited overstaining of their interior by mAb414-AF594 (Fig. S6). These immunostaining results indicate the presence of a nuclear membrane on the surfaces of a certain number of nucleus-like structures observed, thus confirming the successful reconstitution of the nucleus in the GUVs.

### Quantitative characterization of nucleus-like structures in GUVs

To characterize nucleus-like structures containing DNA and GFP-NLS accumulated through nuclear import (top row of Fig. 3A and Fig. S2), we analyzed fluorescence in GUVs formed under each of NLS(+) and NLS(-) conditions and incubated for nuclear assembly reactions. We first defined C1 and C2 compartments in GUVs containing a nucleus-like structure carrying dense DNA (Fig. 4A) and extracted the mean fluorescence intensity in C1 and C2 compartments (Fig. S8). We then computed the ratio of the fluorescence intensity (Fig. 4B-C). The violin plot of SYTO 41 intensity ratio showed similar distributions and mean values of approximately 2.3 under both NLS(+) and NLS(-) conditions (see Table S13 for the result of t-test), in Fig. 4B. In contrast, the distributions of GFP intensity ratio appeared significantly different between the two conditions (*p* = 5.37e-68; see Table S14 for the result of t-test), with mean values of approximately 1.6 for NLS(+) and 1.2 for NLS(-), as seen in Fig. 4C. These results suggest: *i*) the presence of dense DNA in the C1 compartment, and *ii*) significantly higher accumulation of GFP-NLS in the C1 compartment under NLS(+) condition than NLS(-) condition.

**Fig. 4.**
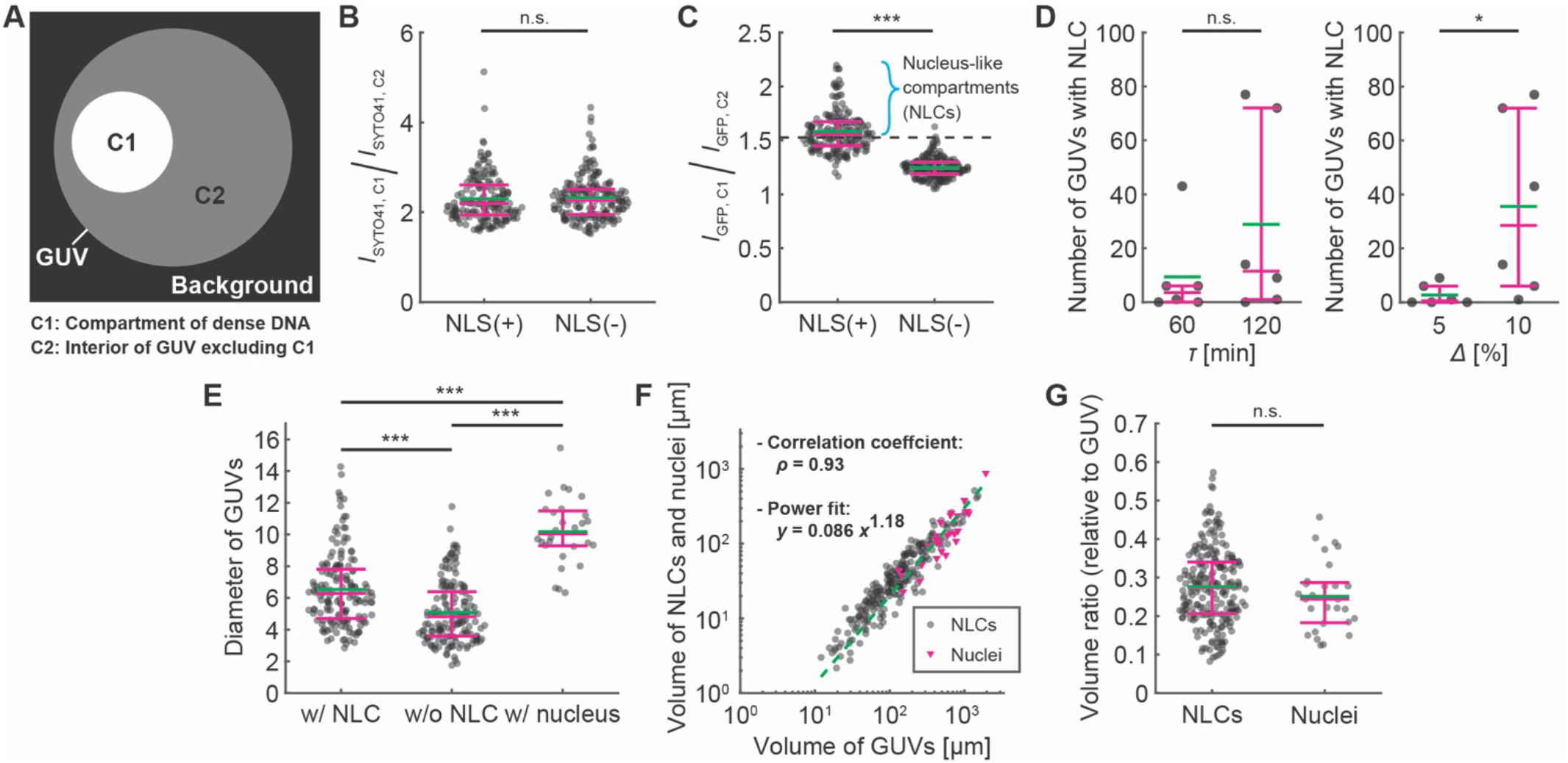
Quantitative analysis of NLCs and nuclei formed in GUVs. (**A**) Definition of internal compartments in a GUV. (**B**) Ratio of SYTO 41 intensity between C1 and C2 compartments in GUVs. *N* = 182 for NLS(+), *N* = 193 for NLS(-). (**C**) Ratio of GFP intensity between C1 and C2 compartments. The threshold of GFP-accumulation is mean + 3SD of NLS(-), indicated with a black dashed line (threshold value: 1.5). (**D**) Number of NLC-containing GUVs in the observed GUVs plotted against *τ* (left) and *Δ* (right). (**E**) Diameter of GUVs at *τ* = 120 min and *Δ* = 10%. GUVs containing an NLC are denoted as “w/ NLC”, GUVs without NLC as “w/o NLC”, and GUVs containing a nucleus as “w/ nucleus”. *N* = 163 for w/ NLC and w/o NLC, *N* =28 for w/ nucleus. (**F**) Volumes of GUVs containing an NLC and that of NLCs are plotted with grey circles. Volumes of GUVs containing a nucleus are plotted in magenta triangles. The green dashed line is a result of power fit (*y* = 0.086 *x*^1.18^). *N* = 229 for NLCs, *N* = 28 for nuclei. (**G**) Ratio of the volume of NLCs and nuclei to the volume of the GUVs. In panels B-E and G, magenta bars indicate the Q1-Q3 quartiles, while a green bar indicates the mean. See “Materials and Methods” for the details on statistical analysis and Tables S13-20 for the results.

To evaluate the impact of waiting time and chloroform addition on the formation of nucleus-like structures in GUVs, we quantified structures containing both dense DNA and accumulated GFP-NLS. “Nucleus-like compartments” (NLCs) were defined as C1 compartments having significantly higher GFP intensity than C2 compartments, determined by a threshold GFP intensity ratio of ∼1.5 (Fig. 4C; see “Materials and Methods” for the details on the definition and identification of NLCs). We observed successful formation of NLC-containing GUVs only under conditions where both waiting time and chloroform volume were non-zero (Fig. 4D). Among these parameters, we confirmed that only chloroform volume had a significant effect on the number of NLC-containing GUVs, with no significant interaction between the two parameters (Tables S15-17 and Fig. 4D). These results indicate that chloroform addition promotes the formation of NLCs in GUVs.

To assess the impact of the GUV size on the successful formation of NLCs and nuclei confirmed by immunostaining, we calculated the diameter of GUVs observed under the condition of *τ* = 120 min and *Δ* = 10%. Statistically significant differences were observed between all combinations of GUVs containing an NLC, GUVs without NLC, and GUVs containing a nucleus as shown in Fig. 4E (see “Materials and Methods” for the details on statistical analysis and Tables S18-19 for the results). The mean diameter of GUVs containing an NLC was 6.6 μm, that of GUVs without NLC was 5.1 μm, and that of GUVs containing a nucleus was 10.2 μm as indicated with green bars in Fig. 4E. In addition, we plotted the diameters of GUVs containing an NLC (grey circles) and a nucleus (magenta triangles) against observed diameters of NLCs and the nuclei and found a positive correlation between GUV volume and NLC and nucleus volume (correlation coefficient: *ρ* ∼ 0.93, power law fit result: *y* = 0.086 *x*^1.18^), as shown in Fig. 4F. The corresponding NLC-to-GUV volume ratio and nucleus-to-GUV volume ratio were approximately 0.27 for NLCs and 0.25 for nuclei, as shown in Fig. 4G (see “Materials and Methods” for the details on statistical analysis and Table S20 for the result). These results suggest three points: *i*) NLC formation and nucleus formation are more likely to occur in larger GUVs within the observed diameter range, *ii*) the size of NLCs and nuclei tends to increase with the size of GUVs within the observed volume range, and *iii*) the volume ratio between NLC and GUV and that between nucleus and GUV have similar distributions.

## Discussion

In this study, we achieved the reconstitution of the cell nucleus in GUVs formed by the inverted emulsion method. Implementation of prolonged waiting time and addition of chloroform into lipid-dispersed oil improved the number of GUVs formed by the inverted emulsion method and the proportion of GUVs encapsulating sperm chromatin-added *Xenopus* interphase egg extract in the observed GUVs (Fig. 2). We found the formation of numerous nucleus-like structures in the GUVs demonstrating their nuclear import capability (Figs. 3A and S2). Moreover, we confirmed the presence of nuclear pores on their surfaces (Figs. 3B and S4), hence the presence of a nuclear membrane, by investigating through immunostaining of nuclear pore complexes. Furthermore, we identified an increase in the assembly frequency of nucleus-like compartments (NLCs), particularly along with an increase of added chloroform volume, as seen in Fig. 4D, and discovered a positive correlation between the size of GUVs and the size of internal NLCs and nuclei (Fig. 4F).

In our investigation, we noted that the implementation of waiting time and the addition of chloroform not only increased the number of GUVs encapsulating the egg extract but also enabled the formation of GUVs encapsulating a high concentration of the egg extract (Fig. S8). We hypothesize that implemented waiting time and added chloroform in lipid-dispersed oil enhance the adsorption of lipid molecules at the oil-water interface and facilitate the formation of GUVs with more stable lipid membranes by the inverted emulsion method (*25,26*). As a result, leakage of the encapsulated egg extract from the interior of the GUVs could have been reduced owing to the increased stability of the GUV membrane, while the internal solution of GUVs formed by the inverted emulsion method normally experiences dilution of the internal content (*29,30*), as demonstrated in Figs. S1 and S8. Provided that the probability of the nuclear assembly event decreases through the loss of required molecules by the extract leakage, the higher concentration of the egg extract in GUVs could have contributed to the observed increase in NLC formation (Fig. 4D).

In general, chloroform is deliberately eliminated from lipid-dispersed oil through evaporation as it is widely considered harmful to biochemical reactions embedded in GUVs (*30-32*). On the other hand, recent investigations employing the inverted emulsion method have demonstrated successful induction of gene expression reactions (*29*) and the bundle formation of cytoskeletal proteins (*33-36*) in GUVs under conditions where chloroform was deliberately added into the lipid-dispersed oil. These studies suggest that chloroform may have limited impact on biochemical reactions in GUVs, at least at the concentrations utilized in these studies. Therefore, we adopted chloroform concentrations in our lipid-dissolved oil (*Δ* = 5/10%) consistent with those used in previous studies, leading to the formation of the numerous egg extract-encapsulating GUVs (Fig. 2) and successful nuclear and NLC assembly in the GUVs (Figs. 3, 4D-G, S2, and S4).

The reconstitution of cell nuclei under micrometer-scale three-dimensional confinement has previously relied on water-in-oil emulsions, where the confirmation of assembled nuclei was achieved only through a nuclear import assay as shown in Fig. S9 (*20-22*). In contrast, our approach involves encapsulating the egg extract in a lipid bilayer of GUVs and reconstitutes a nucleus in the GUVs. The semi-permeability of the lipid bilayer allowed the exchange of molecules between the inside and outside of the GUVs, while essentially isolating the internal solution from the outer environment in confinement. This setup enabled the characterization of nucleus-like structures in an additional approach, similarly to previous studies of *in vitro* nuclei formed in the bulk extract. We first identified the formation of nucleus-like structures in GUVs by a nuclear import assay (Fig. 3A) and additionally conducted their molecular characterization through immunostaining (Fig. 3B). Our findings, indicating that only approximately 17% of NLCs carried nuclear pore complexes on their surfaces, highlight the significance of GUV encapsulation in providing a more comprehensive understanding of *in vitro* nuclei formed under confinement in GUVs.

We obtained several insights into size. First, we observed a significantly higher mean diameter of ∼10.2 μm for GUVs containing a nucleus, as shown in Fig. 4E, compared to the mean diameter of GUVs with or without NLC. We consider that this discrepancy may be attributed to an unidentified effect of the immunostaining procedure developed in this study but not necessarily a prerequisite for nuclear assembly to occur. Furthermore, we confirmed a proportional relationship between the volume of GUVs and that of NLCs and nuclei (Fig. 4F), where the evaluated power-law exponent fitted to 1.18. This exponent is larger than the known exponents (0.7-0.9) between cell volume and nucleus volume in various natural cells (*37*), resulting in a relatively higher nuclear-to-cytoplasmic (N/C) volume ratio in these GUVs compared to many natural cells (Fig. 4G). We attribute this high N/C ratio to the large volumes of sperm chromatin encapsulated in the GUVs. Additionally, we discovered that the size range of GUVs containing an NLC and nucleus spanned approximately 1 to 16 μm, consistent with reported dimensions of GUVs encapsulating *Xenopus* egg extract of unspecified types (*24*). In contrast, we confirmed the presence of larger emulsions in the lipid-dispersed oil just before the phase transfer procedure in the inverted emulsion method (Fig. S10). These observations might be linked to instances of GUV rupture arising from the inverted emulsion method (*31,38,39*). Lastly, we found that a significant subset of the formed GUVs exhibited sizes smaller than the standard dimensions of *Xenopus laevis* sperm chromatin (approximately 10 μm along the long axis), suggesting the potential encapsulation of partial sperm chromatin fragments within the GUVs smaller than 10 μm. Considering the broader applicability of the proposed protocol, it is anticipated that the protocol could facilitate the complete encapsulation of arbitrarily long DNA molecules, including those at a genome-scale. To achieve such encapsulation, it might be necessary to investigate methods for handling long DNA molecules to prevent fragmentation due to physical forces or to develop techniques for sorting GUVs to ensure the complete encapsulation of target DNA.

Taken together, this study has enabled the reconstitution of cell nuclei in GUVs, opening a range of potential applications. Firstly, it may facilitate the investigation of the interaction between the cell cortex and nuclear assembly (*23*), which has likely been hindered by the absence of cell cortex anchoring proteins at the boundary of water-in-oil emulsions (*40*). Moreover, the established protocol holds promise for evolving into a method for transplanting artificial chromosomes into various eukaryotic cells, integrating with artificial chromosome/genome technologies (*41-44*) and GUV-cell fusion techniques (*45,46*). For chromosome transplantation, further optimization of lipid adsorption to the oil-water interface and investigation into phase transfer conditions in the inverted emulsion method might be necessary to form large GUVs encapsulating DNA of arbitrary size. Finally, we believe that the reconstituted cell nuclei in GUVs will serve as a powerful tool for constructing artificial cellular systems closely mimicking eukaryotic cells in bottom-up synthetic biology research.

## Materials and Methods

### Preparation of interphase egg extract and sperm chromatin of the frog *Xenopus laevis*

Interphase egg extract was prepared as described previously (*47*) with modifications as follows: to convert the cell cycle stage of the egg extract from mitosis into interphase, unfertilized eggs were crushed using KMH (100 mM KCl, 2.5 mM MgCl_2_, 20 mM HEPES-KOH at pH 7.7) containing 1 mM CaCl_2_ rather than a calcium-ionophore treatment; to completely separate the egg extract from insoluble material (*e*.*g*., actin filaments and mitochondria), crude egg extract isolated by the initial centrifugation was diluted in KMH (10% volume of the original extract), and then clarified by another round of centrifugation at 45,000×*g*. Sperm chromatin was prepared as described previously (*47*). In every preparation of the egg extract and sperm chromatin, their quality was evaluated by monitoring nuclear assembly in the bulk format (bottom row of Fig. 3B and Fig. S5) and in water-in-oil emulsions (Fig. S9) under a fluorescence microscope.

### Preparation of GFP-NLS

GFP-NLS in Figs. 3B, 4B NLS(+), 4C, 4E w/ nucleus, 4F Nuclei, S1, S4-6, S10 was prepared by following a standard protocol (*48*). GFP-NLS used in other sections of this paper was prepared by Dr. Yuki Hara by following a similar standard protocol. Briefly, *Escherichia coli* BL21 (DE3) cells were transformed with a pGEX vector containing a GFP-NLS gene. The cells were incubated in four 250 mL aliquots of LB medium at 30°C with shaking at 250 RPM until OD_600_ ∼ 0.8. 1 mM IPTG was added, and the cultures were further incubated for 5 h. The cells were harvested by centrifugation, and the pellets were stored at -80°C until next day. On the following day, the frozen cell pellets were resuspended in PBS lysis buffer (consisting of PBS pH 7.2 and 1% Triton X-100) and lysed by gentle sonication. Then, the lysate was centrifuged at 12,000×*g*, 4°C for 15 min. The supernatant was collected and mixed with a 50:50 slurry of glutathione-Sepharose beads in chilled PBS lysis buffer. The bead-cell mix was incubated at 4°C to trap proteins on the beads. Then, the beads were washed with ice-cold PBS containing protease inhibitors and loaded into a disposable chromatography column. Trapped GST-GFP-NLS was eluted with ice-cold 50 mM Tris-HCl (pH 8.0) containing 20 mM reduced glutathione. Molecular weight of the collected GST-GFP-NLS was confirmed by polyacrylamide gel electrophoresis. Throughout this study, GST-GFP-NLS is referred to as “GFP-NLS”.

### Intra-GUV nuclear assembly protocol

To achieve reconstitution of cell nuclei in GUVs, we designed an experimental protocol as illustrated in Fig. 1. Briefly, the protocol consists of three steps: 1) preparation of *Xenopus* interphase egg extract supplemented with sperm chromatin, 2) formation of GUVs encapsulating the sperm chromatin-added interphase egg extract by the inverted emulsion method, and 3) induction of nuclear assembly reactions in the formed GUVs. In the first step, the sperm chromatin-added egg extract was prepared for *in vitro* nuclear assembly reactions (*15-17*) as depicted in Fig. 1A. In the second step, GUVs encapsulating the egg extract were formed at a chilled temperature by the inverted emulsion method (*25*) with two modifications to the conventional protocol: *i*) implementing prolonged waiting time and *ii*) adding chloroform into lipid-dispersed oil to facilitate the formation of GUVs encapsulating the egg extract as illustrated in Fig. 1B. In the third step, the GUVs were incubated at 22-23ºC to initiate the assembly of interphase nuclei within them as depicted in Fig. 1C.

### Formation of GUVs encapsulating *Xenopus* interphase egg extract

The lipid-dispersed oil (lipid-oil) was prepared using the following procedure. 1-palmitoyl-2-oleoyl-glycero-3-phosphocholine (POPC) at a concentration of 25 mg mL^-1^ in chloroform, and 1,2-dioleoyl-sn-glycero-3-phosphoethanolamine-N-(lissamine rhodamine B sulfonyl) ammonium salt (RhPE) at 1 mg mL^-1^ in chloroform, both purchased from Avanti Polar Lipids (USA), were taken to a disposable glass bottle with a screw top (Laboran, As One, Japan). The chloroform was evaporated in a desiccator for 30-60 min using a vacuum pump. Mineral oil (M5904, Sigma Aldrich) was then immediately added to the bottle. The final composition of the lipid-oil was 2.5 mM of POPC with 0.1% (mole percent) RhPE in mineral oil. For the GUVs in immunostaining experiment, RhPE was not added. Subsequently, chloroform (*Δ* = 0/5/10% depending on the experimental condition) was added using a disposable pipette. The screw top was tightly closed, and a strip of adhesive tape was attached to ensure sealing. The lipid-oil was sonicated at 60°C for 1 h and stored at room temperature until use. In all experiments, the lipid-oil was consumed within a week after preparation. The lipid-oil was sonicated at 60°C for 1 h before each use.

GUVs were formed by the inverted emulsion method (*25*) according to the following protocol. First, two 1.5 mL disposable tubes and a 2.0 mL disposable centrifuge tube were prepared. 150 μL of egg lysis buffer (250 mM sucrose, 10 mM HEPES-KOH pH 7.7, 50 mM KCl, 2.5 mM MgCl_2_, and 1 mM dithiothreitol) (*49*) was added to each of the 1.5 mL tubes, and 1.0 mL of lipid-oil was added to the 2.0 mL centrifuge tube. All tubes were then placed on ice. Next, 25 μL of frozen *Xenopus* interphase egg extract stored at -80°C was thawed on ice. Subsequently, 0.5 μL of energy mix (50 mM ATP, 50 mM MgCl_2_, 10 mM phosphocreatine, 100 μg mL^-1^ creatine kinase, pH 7.5 adjusted with NaOH) and 0.5 μL of GFP-NLS (3.4 mg mL^-1^ for experiments in Figs. 2, 3A top, 4D, 4E w/ and w/o NLC, 4F-G NLCs, S2, S8-9, S11-12 and 11.1 mg mL^-1^ for experiments in Figs. 3B, 4B NLS(+), 4C, 4E w/ nucleus, 4F Nuclei, 4G Nuclei, S1, S4-6, S7 NLS(+), S10) for NLS(+) condition or GFP (7.7 mg mL^-1^ for experiments in Figs. 3A bottom, 4B-C NLS(-), S3, S7 NLS(-); purchased from Chromotek, Germany) for NLS(-) condition were added to the thawed egg extract. This mixture was briefly vortexed and spun down. Subsequently, 0.5 μL of thawed *Xenopus* sperm chromatin was added on ice and the extract was mixed by gently pipetting four times. 25 μL of this extract mix was then added to 1 mL of chilled lipid-oil in the 2.0 mL centrifuge tube. Additionally, 150 μL of lipid-oil was layered on top of the egg lysis buffer in the two 1.5 mL tubes on ice. These three tubes were incubated on ice for the designated duration of waiting time (*τ* = 0/60/120 min).

Following the incubation of the oil-water interface on ice, the mixture of lipid-oil and the egg extract was vortexed for 1 min at 3,200 RPM using a Vortex-Genie 2 (Scientific Industries, Inc.). 500 μL of the vortexed mixture of the egg extract in lipid-oil was immediately layered on top of each of the chilled lipid-oil layers in the 1.5 mL tubes. The two 1.5 mL tubes were centrifuged at 9,000×*g*, 4°C for 30 min. During this centrifugation, a new 1.5 mL tube filled with 1 mL of the egg lysis buffer was prepared and chilled on ice. Subsequently, the centrifuged tubes were collected, and the formed GUVs in the bottom pellet were transferred into 1 mL of the chilled egg lysis buffer in a 1.5 mL tube by piercing the side wall of the pellet in the tube with a syringe needle and applying air pressure from the open top of the tube using an index finger. Finally, the collected GUVs dispersed in the egg lysis buffer were vortexed briefly and kept on ice until further experiment.

### Induction of nuclear assembly reactions

Nuclear assembly in GUVs encapsulating sperm chromatin-added *Xenopus* interphase egg extract was conducted by incubating the formed GUVs dispersed in the egg lysis buffer either on a tube rack at 23°C or on a block incubator set at 22°C for 90 min. Note that all GUVs encapsulating the egg extract formed by our inverted emulsion method were incubated for nuclear assembly reactions before fluorescence staining and microscopy.

Nuclear assembly under bulk condition was performed by the following procedure. Firstly, the frozen egg extract, energy mix, GFP-NLS, and sperm chromatin were thawed on ice. Subsequently, 1 μL of energy mix and 1 μL of GFP-NLS were added to 50 μL of the thawed egg extract in a 0.5 mL tube. This egg extract was vortexed briefly, spun down, and placed on ice immediately. Then, 1 μL of sperm chromatin was added to the egg extract. This egg extract was gently pipetted four times, spun down briefly, and placed back on ice immediately. The 0.5 mL tube containing the extract mix was then placed on a block incubator set at 22°C and incubated for 90 min to induce nuclear assembly reactions in the bulk egg extract.

### Fluorescence staining and preparation of glass slide samples for microscopy

GUVs encapsulating the egg extract dispersed in the egg lysis buffer were centrifuged at 200×*g* for 5 min at room temperature. The supernatant was carefully removed, and the sample volume was reduced to 140 μL. For fluorescence staining of DNA, 0.14 μL of 5 mM SYTO 41 (Thermo Fisher Scientific) was added to the sample by a 1,000× dilution. Subsequently, the stained GUVs were briefly vortexed and transferred into the open well of a Frame-Seal chamber (SLF1201, Bio-Rad) attached to a BSA-coated glass slide. The chamber was immediately sealed with a No.1 glass coverslip coated with BSA. To facilitate microscopy, the glass slide sample was immediately inverted so that the No.1 coverslip faced downward, allowing the GUVs to settle towards the bottom surface prior to microscopy.

### Microscopy

An inverted microscope (IX71, Olympus) equipped with an oil immersion objective (PlanApoN 60×, 1.45 NA, Olympus), fluorescence filters (460/80 nm, 520/35 nm, and 617/73 nm Bright-Line single-bandpass filters, Semrock, USA), a laser combiner (405/488/561 nm, ALC5000, Andor Technology, UK), a spinning disk confocal unit (CSU-X1, Yokogawa, Japan), and an EM-CCD camera (iXon X3, Andor Technology, UK) was used for all micrographs in the main text of this paper. Imaging sequences were programmed and executed using iQ2 software (Andor Technology, UK).

### Immunostaining of nuclear pore complexes on nuclear membranes

Immunostaining of nuclear pore complexes (NPCs) on the nuclei formed in the bulk form of the egg extract and in GUVs encapsulating the egg extract was conducted as follows.

For fixing nuclei formed in bulk form of the egg extract, 50 μL of the nuclei-containing egg extract was first mixed with 150 μL of the egg lysis buffer and pipetted gently. 200 μL of fixation solution (4% glutaraldehyde in 80 mM KCl and 10 mM Tris-HCl pH 7.7) was added, and the tube was immediately placed on ice and incubated for 1 h. After the incubation, the fixed nuclei were transferred to a 1.5 mL tube filled with 1 mL of quenching solution (10 mM Tris-HCl pH 7.7, 250 mM sucrose, 50 mM KCl, 1% BSA) on ice. The sample was centrifuged at 200×*g*, 4°C for 5 min, and the supernatant was gently removed. The sample was then centrifuged again, and the supernatant was removed down to 100 μL. To stain NPCs on the membrane of the fixed nuclei, 1 μL of 0.5 mg mL^-1^ mAb414-AF594 was added and gently mixed by pipetting. The fixed nuclei with mAb414-AF594 were incubated on ice overnight. On the next day, the sample was centrifuged at 200×*g*, 4°C for 5 min. The supernatant was removed down to 100 μL, and 0.1 μL of 5 mM SYTO 41 was added for DNA staining. The entire volume of the immunostained nuclei was enclosed in a Frame-Seal chamber between a No.1 glass coverslip and a glass slide, both of which were coated with BSA.

The fixation of nuclei formed in GUVs encapsulating the egg extract was performed in the following procedure. The GUVs were centrifuged at 200×*g*, 4ºC for 5 min, and the supernatant was removed down to 100 μL. The tube containing the GUVs was placed on ice, and 100 μL of the fixation solution was added. After 1 h of incubation on ice, the fixed GUVs were mixed with 1 mL of quenching solution and centrifuged at 1,000×*g* for 5 min at 4ºC. The supernatant was removed, and the volume was reduced to 100 μL. The quenching and washing of the fixed GUVs were repeated once, and the sample volume was adjusted to 100 μL. The fixed GUVs with NPC-antibodies were then incubated on ice overnight. The following day, 1 mL of quenching solution was added, the sample was centrifuged at 1,000×*g*, 4ºC for 5 min, and the supernatant was reduced down to 100 μL. 0.1 μL of 5 mM SYTO 41 was added for DNA staining, and the immunostained GUVs were enclosed in a Frame-Seal chamber between a No.1 glass coverslip and a glass slide, both of which were coated with BSA.

### Image segmentation of GUVs

Image segmentation of GUVs on acquired micrographs was performed using *cellpose 2*.*0* (*50,51*) implemented in Python. The envelopes of GUVs were primarily segmented using a built-in model (“CP”), and the quality of the segments was manually inspected and corrected on the graphical user interface (GUI) of *cellpose 2*.*0*.

### Number, internal GFP intensity, and diameter of GUVs

The analysis of GUV fluorescence using above generated segment masks was performed in MATLAB. The number of GUVs was determined by computing the number of segments. The mean GFP intensity in the GUVs, denoted as *I*_GFP_, was computed by extracting intensity values in the GFP channel using the segmentation mask and taking their mean. The diameter of the GUVs was defined as the reduced diameter of each circle having a corresponding area of individual segments. To determine the mean GFP intensity in the background of each micrograph, three arbitrary rectangular regions with at least 1,056 pixels were carefully selected in the background of each micrograph. The corresponding mean GFP fluorescence intensity was computed for each rectangular region, and the average of the three means was calculated. The threshold of GFP encapsulation (dashed lines in Fig. S8) was defined by mean + 3SD of the mean GFP intensity in each background.

### Exclusion of GUVs containing aggregated GFP

GUVs containing aggregated GFP were manually excluded from the dataset used to compute the proportion of GUVs encapsulating the GFP-dispersed egg extract in the observed GUVs (Fig. 2C). This exclusion aimed to mitigate artifacts in the computed proportion arising from GFP aggregation in three-dimensional space. This issue was observed in all micrographs acquired after completing the entire protocol of nuclear assembly in GUVs (Fig. 1). Since the GUVs in the micrographs of Fig. 2 had already undergone 23ºC incubation for nuclear assembly reactions, several GUVs in the acquired micrographs exhibited positive indications of nuclear import by GFP-NLS within them. Theoretically, the mean GFP intensity computed from a confocal micrograph of such GUVs with aggregated GFP always yielded a higher mean GFP intensity than a confocal micrograph of GUVs without GFP aggregation even if the same amount of GFP was encapsulated. Consequently, the mean GFP intensity of the GUVs containing the aggregation did not precisely reflect the concentration of GFP encapsulated in the GUV. Because the purpose of the analysis in Fig. 2C was to evaluate GFP encapsulation in the GUVs, we considered the inclusion of data from GUVs carrying such aggregated GFP inadequate and excluded the GUVs from this specific analysis in Fig. 2C. The same parameter computed from data including the GUVs containing aggregated GFP is shown in Fig. S11. Additionally, raw GFP intensity values in the GUV lumen and the background of these GUVs are plotted in Fig. S12. Comparisons between plots excluding the GUVs containing aggregated GFP (Figs. 2C and S8) and those including them (Figs. S11-12) indicate only minor changes.

### Definition and identification of NLCs in GUVs

To differentiate between C1 compartments (Fig. 4A) containing a significantly high level of GFP-NLS from those with a lower level, we set a threshold for the GFP intensity ratio, as indicated by the dashed line in Fig. 4C. This threshold was determined as the mean + 3SD of the GFP intensity ratio under NLS(-) conditions (dashed line in Fig. 4C). We used this threshold to interpret accumulation of GFP-NLS in C1 compartment: if the ratio exceeded the threshold, it indicated GFP-NLS accumulation. Utilizing this threshold, we defined a “nucleus-like compartment” (NLC) as a C1 compartment with a GFP intensity ratio higher than the threshold.

To identify NLCs in GUVs, GUVs containing dense DNA (C1 compartment in Fig. 4A) were initially identified in micrographs through manual inspection. Subsequently, the envelopes of the GUVs and their internal C1 compartments were manually segmented in RhPE and SYTO41 channels using the interactive *drawfreehand* function in MATLAB. Mean intensity values of SYTO 41 signals in the C1 and C2 compartments were then computed using the segment masks. Similarly, mean GFP intensities in the C1 and C2 compartments were computed using the same segment masks. Next, the ratio of mean intensities in the C1 and C2 compartments was computed for both the SYTO 41 and GFP channels. The threshold for the GFP intensity ratio was determined based on the above definition (computed threshold: ∼1.5). Furthermore, to ensure the encapsulation of DNA and GFP-NLS in the NLCs, the mean fluorescence intensity of both SYTO 41 and GFP in the background (Fig. S7) was calculated. This computation was done by extracting three arbitrary rectangular regions from the background, each containing at least 1,517 pixels. The mean fluorescence intensity of each region was computed, and the average of these three mean values was taken.

### Number and size of GUVs containing an NLC

The number of GUVs containing an NLC was determined by manually inspecting all confocal micrographs of GUVs encapsulating the egg extract after 90 minutes of incubation at 22°C. GUVs carrying dense DNA (C1 compartment) were manually identified, and each C1 compartment was evaluated to determine whether it contained an NLC, following the established definition. The diameters of GUVs containing an NLC and those not containing an NLC at *τ* = 120 minutes and *Δ* = 10%, as shown in Fig. 4E, were computed using the following procedure. Accurate manual image segmentation of GUVs containing an NLC was conducted when all NLCs were identified. The same number of segments from GUVs without an NLC were randomly selected from the micrographs. Using the segments of these GUVs, the reduced diameter was computed from their segment area. The volumes of GUVs containing an NLC and nucleus, as well as the volumes of the corresponding NLC and nucleus, were computed from the segment area by calculating the reduced radius and then the corresponding reduced volume as a sphere.

### Statistical analysis

In the statistical analyses depicted in Figs. 2B-C, 2E, 2F, 4D, and S11, we conducted a two-way ANOVA followed by *post-hoc* Tukey’s HSD tests, focusing either on the duration of waiting time or the added volume of chloroform. For the statistical analyses in Figs. 4B-C, we performed a two-sample t-test. For the statistical analysis in Fig. 4E, we used the Kruskal-Wallis test followed by a *post-hoc* Mann-Whitney U test with Bonferroni’s correction. In the statistical analysis shown in Fig. 4G, we employed the Wilcoxon rank-sum test. For all analyses, resultant *p*-values were rounded to the first significant digit and annotated using the conventional method: “n.s.” (not significant) for *p* > 0.05, “*” for *p* ≤ 0.05, “**” for *p* ≤ 0.01, and “***” for *p* ≤ 0.001. The results of all statistical analyses are summarized in Tables S1-23.

## Acknowledgments

We thank Dr Yuki Hara for kindly offering purified GFP-NLS and the corresponding plasmid.

## Author contributions

Conceptualization: ST1(STakeuchi), MO, ST2(STakamori)

Methodology: ST1, MO, KS, TO, ST2, HM, TK, MS

Investigation: KS, ST2, MS

Visualization: ST2

Supervision: ST1, MO, KS, TO, HM, TK, MS

Writing—original draft: ST2

Writing—review & editing: ST1, MO, KS, TO, ST2, HM, TK, MS

## Data and materials availability

All data needed to evaluate the conclusions in the paper are available in the main text or the Supplementary Materials.

## Supplementary Materials

Please see a separately submitted ‘Supplementary Materials’ for supplementary information: Supplementary Text, Figs. S1 to S12, Tables S1 to S23.

## References

(1) H. Jia, P. Schwille, Bottom-up synthetic biology: reconstitution in space and time. Current opinion in biotechnology 60, 179–187 (2019).

(2) F. C. Simmel, Synthetic organelles. Emerging Topics in Life Sciences 3(5), 587–595 (2019).

(3) B. C. Buddingh’, J. C. van Hest, Artificial cells: synthetic compartments with life-like functionality and adaptivity. Accounts of chemical research 50(4), 769–777 (2017).

(4) N. J. Gaut, K. P. Adamala, Reconstituting natural cell elements in synthetic cells. Advanced Biology 5(3), 2000188 (2021).

(5) Y. Elani, Interfacing living and synthetic cells as an emerging frontier in synthetic biology. Angewandte Chemie 133(11), 5662–5671 (2021).

(6) K. Y. Lee, S. J. Park, K. A. Lee, S. H. Kim, H. Kim, Y. Meroz, L. Mahadevan, K. H. Jung, T. K. Ahn, K. K. Parker, K. Shin, Photosynthetic artificial organelles sustain and control ATP-dependent reactions in a protocellular system. Nature biotechnology 36(6), 530–535 (2018).

(7) S. Berhanu, T. Ueda, Y. Kuruma, Artificial photosynthetic cell producing energy for protein synthesis. Nature communications 10(1), 1325 (2019).

(8) B. P. Kumar, J. Fothergill, J. Bretherton, L. Tian, A. J. Patil, S. A. Davis, S. Mann, Chloroplast-containing coacervate micro-droplets as a step towards photosynthetically active membrane-free protocells. Chemical communications 54(29), 3594–3597 (2018).

(9) L. Otrin, N. Marušič, C. Bednarz, T. Vidakovic-Koch, I. Lieberwirth, K. Landfester, K. Sundmacher, Toward artificial mitochondrion: mimicking oxidative phosphorylation in polymer and hybrid membranes. Nano letters 17(11), 6816–6821 (2017).

(10) N. Marušič, L. Otrin, Z. Zhao, R. B. Lira, F. L. Kyrilis, F. Hamdi, P. L. Kastritis, T. Vidaković-Koch, I. Ivanov, K. Sundmacher, R. Dimova, Constructing artificial respiratory chain in polymer compartments: Insights into the interplay between bo3 oxidase and the membrane. Proceedings of the National Academy of Sciences 117(26), 15006–15017 (2020).

(11) N. Westensee, E. Brodszkij, X. Qian, T. F. Marcelino, K. Lefkimmiatis, B. Städler, Mitochondria encapsulation in hydrogel-based artificial cells as ATP producing subunits. Small 17(24), 2007959 (2021).

(12) N. N. Deng, M. Yelleswarapu, L. Zheng, W. T. Huck, Microfluidic assembly of monodisperse vesosomes as artificial cell models. Journal of the American Chemical Society 139(2), 587–590 (2017).

(13) L. Aufinger, F. C. Simmel, Artificial gel-based organelles for spatial organization of cell-free gene expression reactions. Angewandte Chemie International Edition 57(52), 17245–17248 (2018).

(14) K. Göpfrich, I. Platzman, J. P. Spatz, Mastering complexity: towards bottom-up construction of multifunctional eukaryotic synthetic cells. Trends in biotechnology 36(9), 938–951 (2018).

(15) M. J. Lohka, Y. Masui, Formation in vitro of sperm pronuclei and mitotic chromosomes induced by amphibian ooplasmic components. Science 220(4598), 719–721 (1983).

(16) D. D. Newmeyer, J. M. Lucocq, T. R. Bürglin, E. M. De Robertis, Assembly in vitro of nuclei active in nuclear protein transport: ATP is required for nucleoplasmin accumulation. The EMBO journal 5(3), 501–510 (1986).

(17) J. Newport, Nuclear reconstitution in vitro: stages of assembly around protein-free DNA. Cell 48(2), 205–217 (1987).

(18) Y. Hara, C. A. Merten, Dynein-based accumulation of membranes regulates nuclear expansion in Xenopus laevis egg extracts. Developmental cell 33(5), 562–575 (2015).

(19) J. Bisht, P. LeValley, B. Noren, R. McBride, P. Kharkar, A. Kloxin, J. Gatlin, J. Oakey, Light-inducible activation of cell cycle progression in Xenopus egg extracts under microfluidic confinement. Lab on a Chip 19(20), 3499–3511 (2019).

(20) M. C. Good, Encapsulation of Xenopus egg and embryo extract spindle assembly reactions in synthetic cell-like compartments with tunable size. The Mitotic Spindle: Methods and Protocols 87-108 (2016).

(21) Y. Guan, Z. Li, S. Wang, P. M. Barnes, X. Liu, H. Xu, M. Jin, A. P. Liu, Q. Yang, A robust and tunable mitotic oscillator in artificial cells. Elife 7, e33549 (2018).

(22) Y. Guan, S. Wang, M. Jin, H. Xu, Q. Yang, Reconstitution of cell-cycle oscillations in microemulsions of cell-free Xenopus egg extracts. JoVE (Journal of Visualized Experiments) (139), e58240 (2018).

(23) J. G. Bermudez, H. Chen, L. C. Einstein, M. C. Good, Probing the biology of cell boundary conditions through confinement of Xenopus cell-free cytoplasmic extracts. genesis 55(1-2), e23013 (2017).

(24) M. Chiba, M. Miyazaki, S. I. Ishiwata, Quantitative analysis of the lamellarity of giant liposomes prepared by the inverted emulsion method. Biophysical journal, 107(2), 346–354 (2014).

(25) S. Pautot, B. J. Frisken, D. A. Weitz, Production of unilamellar vesicles using an inverted emulsion. Langmuir 19(7), 2870–2879 (2003).

(26) C. Claudet, M. In, G. Massiera, Method to disperse lipids as aggregates in oil for bilayers production. The European Physical Journal E 39, 1–6 (2016).

(27) M. R. Dilsaver, P. Chen, T. A. Thompson, T. Reusser, R. N. Mukherjee, J. Oakey, D. L. Levy, Emerin induces nuclear breakage in Xenopus extract and early embryos. Molecular Biology of the Cell 29(26), 3155–3167 (2018).

(28) M. S. Nord, C. Bernis, S. Carmona, D. C. Garland, A. Travesa, D. J. Forbes, Exportins can inhibit major mitotic assembly events in vitro: membrane fusion, nuclear pore formation, and spindle assembly. Nucleus 11(1), 178–193 (2020).

(29) L. Van de Cauter, F. Fanalista, L. Van Buren, N. De Franceschi, E. Godino, S. Bouw, C. Danelon, C. Dekker, G. H. Koenderink, K. A. Ganzinger, Optimized cDICE for efficient reconstitution of biological systems in giant unilamellar vesicles. ACS Synthetic Biology 10(7), 1690–1702 (2021).

(30) Y. Shimane, Y. Kuruma, Rapid and facile preparation of giant vesicles by the droplet transfer method for artificial cell construction. Frontiers in Bioengineering and Biotechnology 10, 873854 (2022).

(31) K. Nishimura, T. Matsuura, K. Nishimura, T. Sunami, H. Suzuki, T. Yomo, Cell-free protein synthesis inside giant unilamellar vesicles analyzed by flow cytometry. Langmuir 28(22), 8426–8432 (2012).

(32) M. Walczak, R. A. Brady, L. Mancini, C. Contini, R. Rubio-Sánchez, W. T. Kaufhold, P. Cicuta, L. Di Michele, Responsive core-shell DNA particles trigger lipid-membrane disruption and bacteria entrapment. Nature Communications 12(1), 4743 (2021).

(33) E. Loiseau, J. A. Schneider, F. C. Keber, C. Pelzl, G. Massiera, G. Salbreux, A. R. Bausch, Shape remodeling and blebbing of active cytoskeletal vesicles. Science Advances 2(4), e1500465 (2016).

(34) T. Litschel, B. Ramm, R. Maas, M. Heymann, P. Schwille, Beating vesicles: encapsulated protein oscillations cause dynamic membrane deformations. Angewandte Chemie International Edition 57(50), 16286–16290 (2018).

(35) T. Litschel, C. F. Kelley, D. Holz, M. Adeli Koudehi, S. K. Vogel, L. Burbaum, N. Mizuno, D. Vavylonis, P. Schwille, Reconstitution of contractile actomyosin rings in vesicles. Nature communications 12(1), 2254 (2021).

(36) L. Baldauf, F. Frey, M. A. Perez, T. Idema, G. H. Koenderink, Branched actin cortices reconstituted in vesicles sense membrane curvature. Biophysical journal 122(11), 2311–2324 (2023).

(37) Y. Hara, Specialization of nuclear membrane in eukaryotes. Journal of Cell Science 133(12), jcs241869 (2020).

(38) N. Yandrapalli, T. Seemann, T. Robinson, On-chip inverted emulsion method for fast giant vesicle production, handling, and analysis. Micromachines 11(3), 285 (2020).

(39) Moga, N. Yandrapalli, R. Dimova, T. Robinson, Optimization of the inverted emulsion method for high-yield production of biomimetic giant unilamellar vesicles. ChemBioChem 20(20), 2674–2682 (2019).

(40) R. Sakamoto, M. Tanabe, T. Hiraiwa, K. Suzuki, S. I. Ishiwata, Y. T. Maeda, M. Miyazaki, Tug-of-war between actomyosin-driven antagonistic forces determines the positioning symmetry in cell-sized confinement. Nature communications 11(1), 3063 (2020).

(41) W. Murray, J. W. Szostak, Construction of artificial chromosomes in yeast. Nature 305(5931), 189–193 (1983).

(42) J. J. Harrington, G. V. Bokkelen, R. W. Mays, K. Gustashaw, H. F. Willard, Formation of de novo centromeres and construction of first-generation human artificial microchromosomes. Nature genetics 15(4), 345–355 (1997).

(43) D. G. Gibson, G. A. Benders, C. Andrews-Pfannkoch, E. A. Denisova, H. Baden-Tillson, J. Zaveri, T. B. Stockwell, A. Brownley, D. W. Thomas, M. A. Algire, C. Merryman, L. Young, V. N. Noskov, J. I. Glass, J. C. Venter, C. A. Hutchison III, H. O. Smith, Complete chemical synthesis, assembly, and cloning of a Mycoplasma genitalium genome. Science 319(5867), 1215–1220 (2008).

(44) C. A. Hutchison III, R. Y. Chuang, V. N. Noskov, N. Assad-Garcia, T. J. Deerinck, M. H. Ellisman, J. F. Pelletier, Z. Qi, R. A. Richter, E. A. Strychalski, L. Sun, Y. Suzuki, B. Tsvetanova, K. S. Wise, H. O. Smith, J. I. Glass, C. Merryman, D. G. Gibson, J. C. Venter, Design and synthesis of a minimal bacterial genome. Science 351(6280), aad6253 (2016).

(45) C. Saito, T. Ogura, K. Fujiwara, S. Murata, S. I. M. Nomura, Introducing micrometer-sized artificial objects into live cells: A method for cell–giant unilamellar vesicle electrofusion. PLoS One 9(9), e106853 (2014).

(46) S. Takamori, P. Cicuta, S. Takeuchi, L. Di Michele, DNA-assisted selective electrofusion (DASE) of Escherichia coli and giant lipid vesicles. Nanoscale 14(38), 14255–14267 (2022).

(47) K. Shintomi, T. Hirano, A sister chromatid cohesion assay using Xenopus egg extracts. Methods in Molecular Biology 1515, 3–21 (2017).

(48) M. B. Einarson, E. N. Pugacheva, J. R. Orlinick, Preparation of GST fusion proteins. Cold Spring Harbor Protocols 2007(4), pdb-prot4738 (2007).

(49) X. Cheng, J. E. Ferrell Jr, Spontaneous emergence of cell-like organization in Xenopus egg extracts. Science 366(6465), 631–637 (2019).

(50) C. Stringer, T. Wang, M. Michaelos, M. Pachitariu, Cellpose: a generalist algorithm for cellular segmentation. Nature methods 18(1), 100–106 (2021).

(51) M. Pachitariu, C. Stringer, Cellpose 2.0: how to train your own model. Nature methods 19(12), 1634–1641 (2022).

