## Supplementary Materials for "Nuclear assembly in giant unilamellar vesicles encapsulating *Xenopus* egg extract"

Sho Takamori *et al.*

**This PDF file includes:**

Supplementary Text
Figs. S1 to S12
Tables S1 to S23

### Supplementary Text

Formation of GUVs encapsulating a standard dilute solution

GUVs shown in Fig. S1 were formed using the same procedure as described in the main text, except that the inside solution consisted of 350 mM sucrose with 10 mM HEPES-KOH (pH 7.7) and 0.2 mg $mL^{-1}$ GFP, while the outside solution comprised 350 mM glucose with 10 mM HEPES-KOH (pH 7.7). No waiting time was introduced ($\tau$ = 0 min) and no chloroform was added ($\Delta$ = 0 %). The procedures for the preparation of glass slide samples were the same as those described in the main text. Microscopy was also performed in the same procedure described in the main text.

Nuclear assembly in water-in-oil emulsions of the egg extract

Nuclear assembly in water-in-oil emulsions of the egg extract was conducted using the following procedure. The thawed interphase egg extract (50 μL) was combined with energy mix (1 μL, composition detailed in the main text), GFP-NLS (1 μL, composition detailed in the main text), and sperm chromatin (1 μL, composition detailed in the main text) on ice. A 2.0 mL tube containing 1 mL of HFE-7500 (Novec 7500, 3M) with 0.2% Fluo-Surf (The Dolomite Centre Ltd) was chilled on ice. Subsequently, 25 μL of the egg extract was added to the tube. The tube was then vortexed for 1 min at 3,200 RPM using Vortex-Genie 2 (Scientific Industries, Inc.) to generate stable emulsions. These emulsions were placed on a tube rack and incubated for 90 min at 23°C to induce nuclear assembly. After the incubation period, the emulsions in HFE-7500 with 0.2% Fluo-Surf were transferred into a glass slide chamber made of Frame-Seal (SLF1201, Bio-Rad) and sealed with a No.1 glass coverslip. Microscopy was also performed in the same procedure described in the main text.

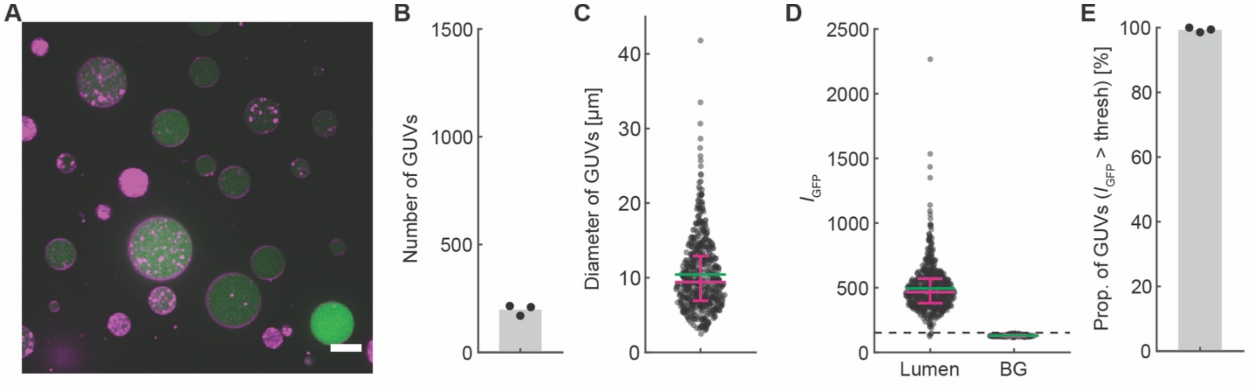


**Fig. S1. GUVs encapsulating a GFP-dispersed solution formed by the inverted emulsion method at $\tau$ = 0 min and $\Delta$ = 0%.**

(**A**) Representative confocal micrograph of GUVs. Magenta indicates RhPE (lipid membrane) and green indicates GFP dispersed in the inside solution of GUVs. Scale bar: 10 µm. (**B**) Number of GUVs. Each point represents the sum of 10 micrographs ($N$ = 1). (**C**) Diameter of GUVs. Sum of 30 micrographs ($N$ = 1). (**D**) Mean GFP intensity in the lumen of GUVs and the background. Sum of 30 micrographs ($N$ = 1). The dashed line represents the mean + 3SD of the background (BG), defined as the threshold of GFP encapsulation in GUVs. (**E**) Proportion of GUVs whose internal mean GFP intensity is higher than the background threshold, indicating the proportion of GUVs encapsulating GFP-dispersed sucrose buffer in the observed GUVs. 10 micrographs per plot ($N$ = 1). In panels C-D, magenta bars indicate Q1-Q3 quartiles and green bars indicate the mean. See Supplementary Text for the details of the experiment.

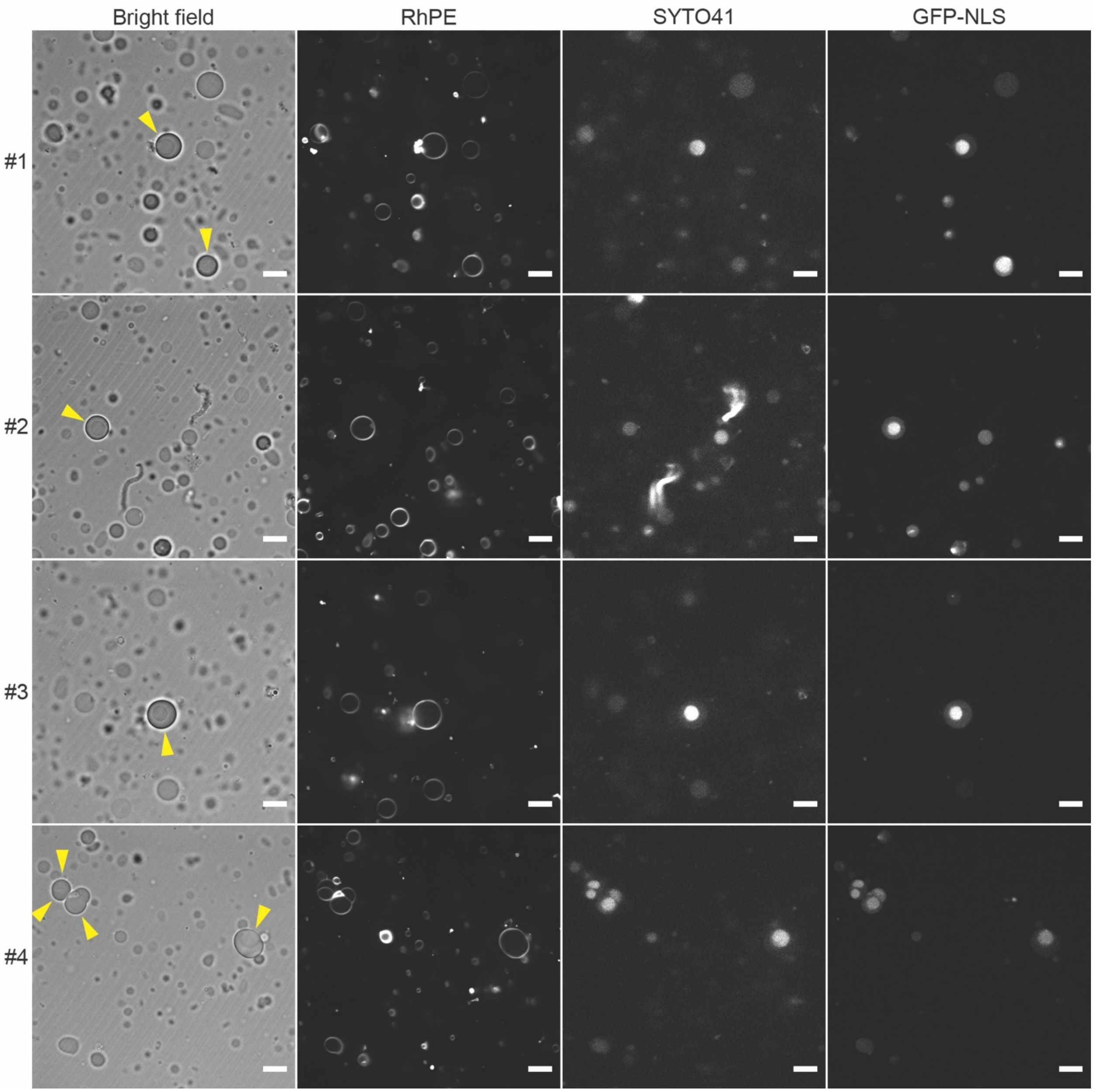


**Fig. S2. Four examples of GUVs with a nucleus-like structure in NLS(+) samples.**
RhPE (18:1 Liss Rhod PE) indicates the lipid membrane. SYTO 41 indicates nucleic acids (mainly DNA). GFP-NLS represents GFP fused with nuclear localization signals (NLS) and dispersed in the egg extract prior to encapsulation into GUVs. GUVs containing a nucleus-like structure are highlighted with yellow arrowheads on the bright field micrographs. Scale bar: 10 µm.

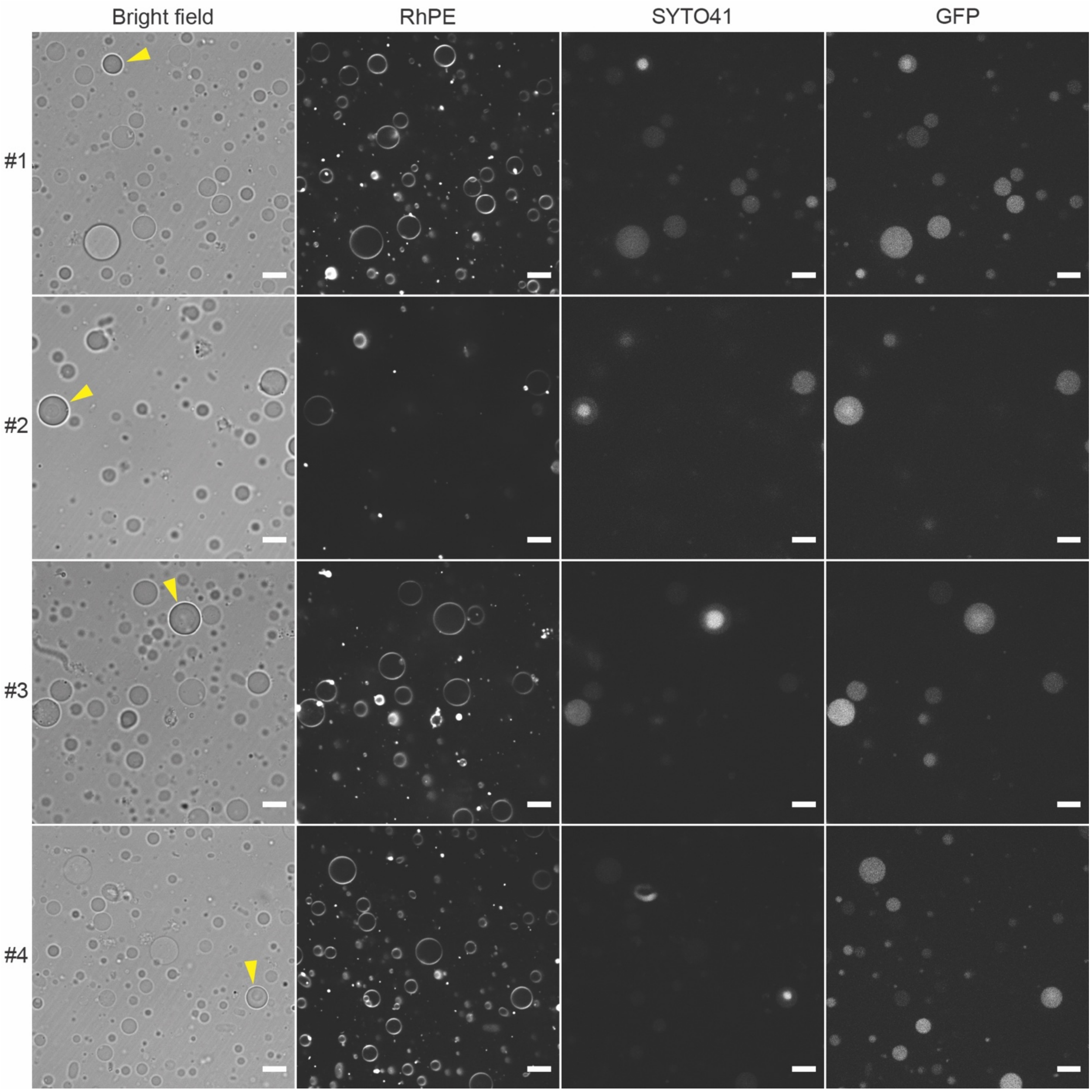


**Fig. S3. Four examples of GUVs with a nucleus-like structure in NLS(-) samples.**
RhPE indicates the lipid membrane. SYTO 41 indicates nucleic acids (mainly DNA). GFP is dispersed in the egg extract prior to encapsulation into GUVs. GUVs containing dense DNA are highlighted with yellow arrowheads on the bright field micrographs. Scale bar: 10 μm.

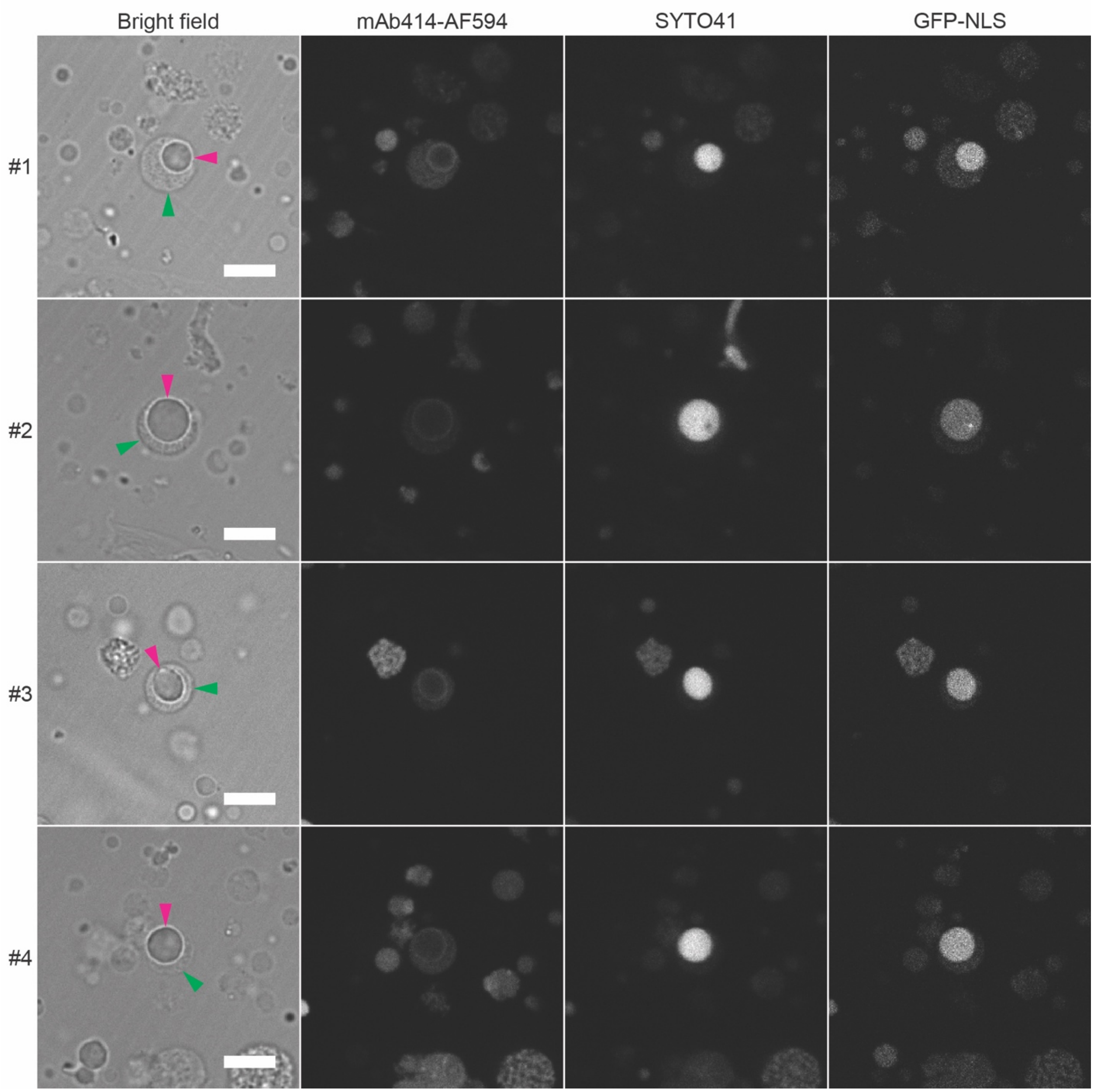


**Fig. S4. Four examples of immunostaining in GUVs containing a nucleus-like structure.** mAb414-AF594 is a monoclonal antibody of nuclear pore complexes conjugated with a fluorophore (Alexa Fluor 594). SYTO 41 indicates nucleic acids (mainly DNA). GFP-NLS is GFP fused with nuclear localization signals (NLS) dispersed in the egg extract prior to the encapsulation into GUVs. See "Materials and Methods" in the main text for the detailed procedure of immunostaining. Arrowheads indicate GUV membrane (green) and nucleus-like structure (magenta). Scale bar: 10 μm.

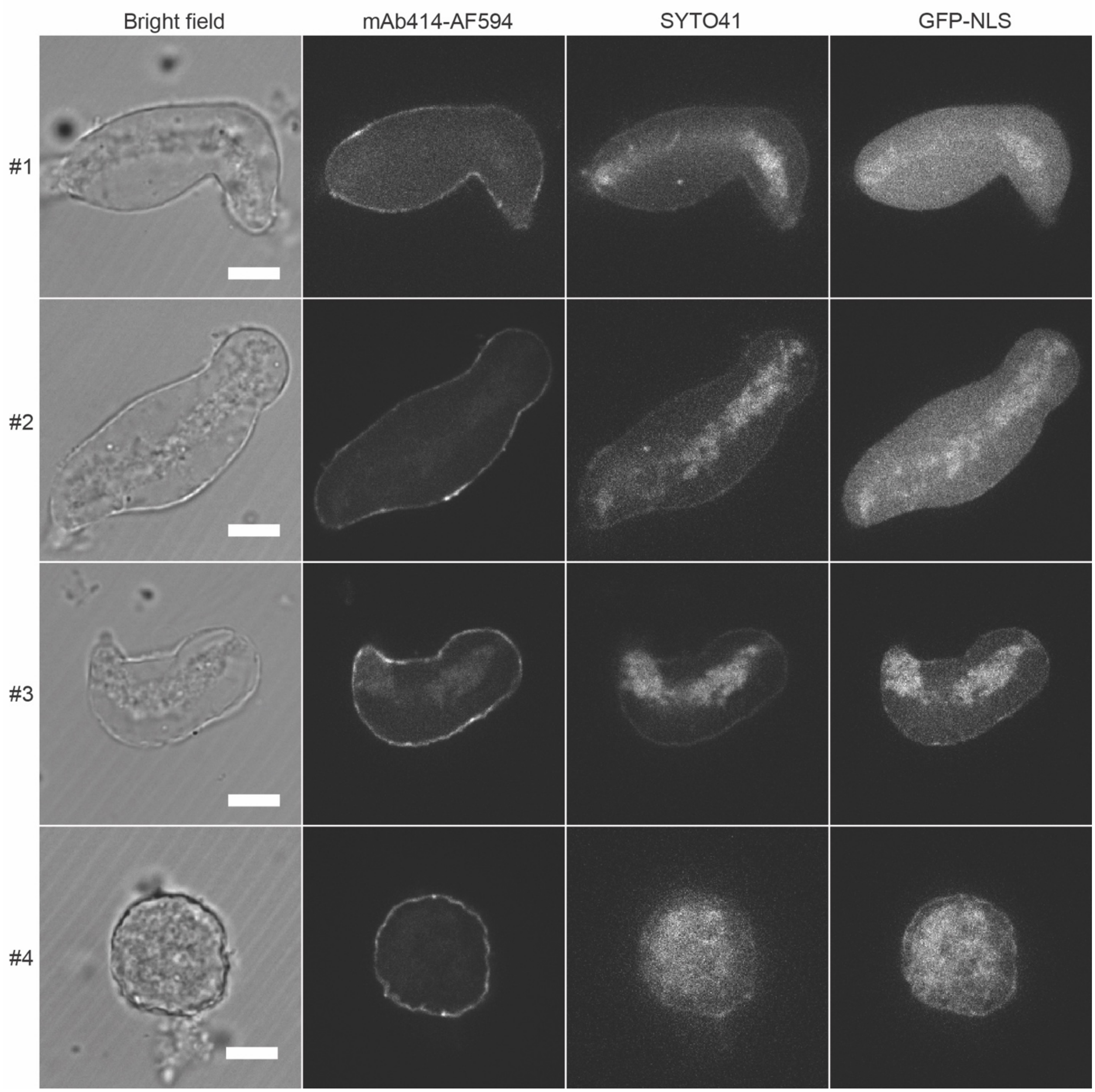


**Fig. S5. Four examples of immunostaining on nuclei formed in bulk form of the egg extract.** mAb414-AF594 is a monoclonal antibody of nuclear pore complexes conjugated with a fluorophore (Alexa Fluor 594). SYTO 41 indicates nucleic acids (mainly DNA). GFP-NLS is GFP fused with nuclear localization signals (NLS) dispersed in the egg extract before nuclear assembly. See "Materials and Methods" in the main text for the detailed procedure of immunostaining. Scale bar: 10 μm.

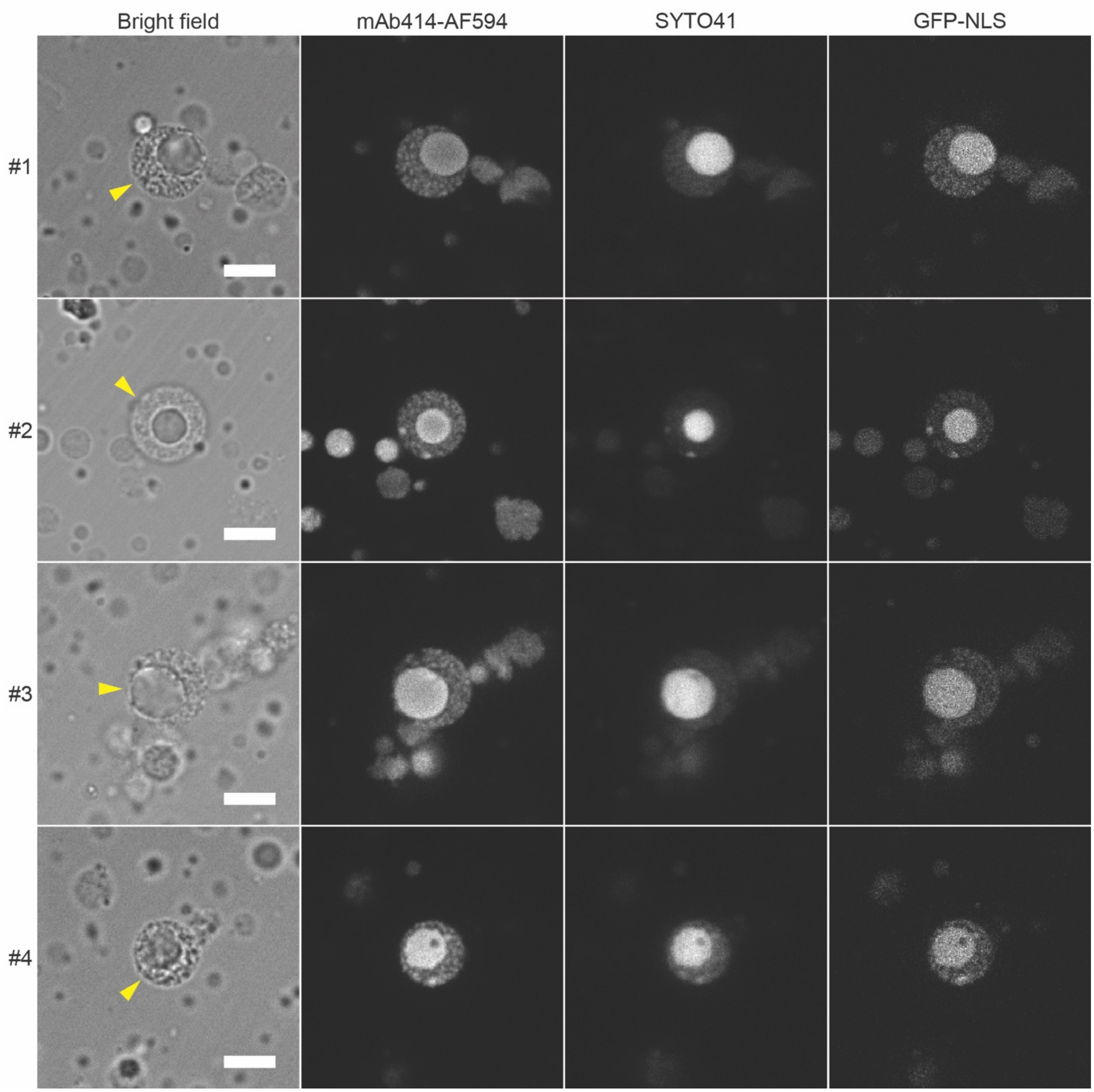


**Fig. S6. Four examples of overstaining in immunostained GUVs containing a nucleus-like structure.**

mAb414-AF594 is a monoclonal antibody of nuclear pore complexes conjugated with a fluorophore (Alexa Fluor 594). SYTO 41 indicates nucleic acids (mainly DNA). GFP-NLS is GFP fused with nuclear localization signals (NLS) dispersed in the egg extract prior to encapsulation. See "Materials and Methods" in the main text for the detailed procedure of immunostaining. GUVs containing a nucleus-like structure are highlighted with yellow arrowheads. Scale bar: 10 µm.

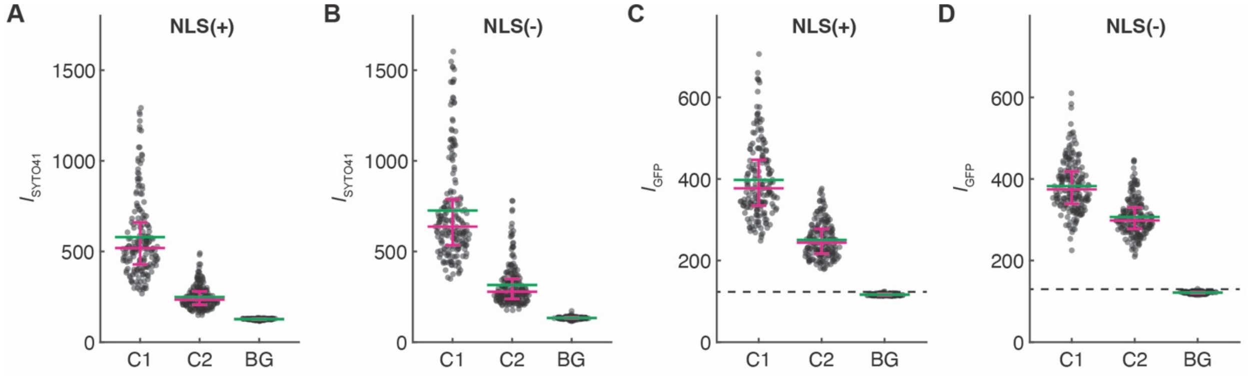


**Fig. S7. Mean fluorescence intensity in GUVs containing a nucleus-like structure.**
(**A**) Mean SYTO 41 intensity in GUVs formed under NLS(+) condition. (**B**) Mean SYTO 41 intensity in GUVs formed under NLS(-) condition. (**C**) Mean GFP intensity in GUVs formed under NLS(+) condition. (**D**) Mean GFP intensity in GUVs formed under NLS(-) condition. "C1" represents a compartment of dense DNA. "C2" represents GUV interior excluding C1 compartment. "BG" is the background. See Fig. 4A in the main text for the graphical definition of C1 and C2 compartments. Magenta bars indicate Q1-Q3 quartiles and green bars indicate the mean. The dashed line indicates the threshold of GFP encapsulation in the GUVs, defined by mean + 3SD of the background (BG).

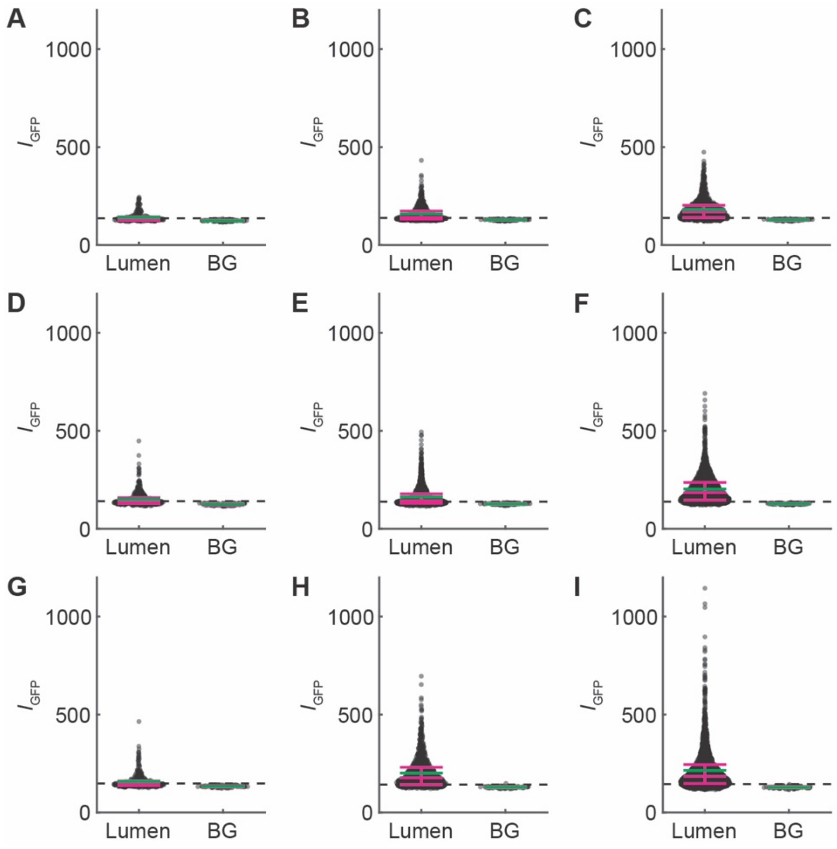


**Fig. S8. Mean GFP intensity in GUVs formed under various conditions of waiting time and chloroform.**

(**A**) $\tau$ = 0 min, $\Delta$ = 0%. (**B**) $\tau$ = 0 min, $\Delta$ = 5%. (**C**) $\tau$ = 0 min, $\Delta$ = 10%. (**D**) $\tau$ = 60 min, $\Delta$ = 0%. (**E**) $\tau$ = 60 min, $\Delta$ = 5%. (**F**) $\tau$ = 60 min, $\Delta$ = 10%. (**G**) $\tau$ = 120 min, $\Delta$ = 0%. (**H**) $\tau$ = 120 min, $\Delta$ = 5%. (**I**) $\tau$ = 120 min, $\Delta$ = 10%. "Lumen" denotes the internal mean GFP intensity, $I_{GFP}$, of GUVs. "BG" denotes the mean GFP intensity of the background at each condition. Magenta bars indicate Q1-Q3 quartiles and green bars indicate the mean. The dashed line indicates the threshold of GFP encapsulation, defined as the mean + 3SD of each background. See "Materials and Methods" in the main text for the details on parameter computation.

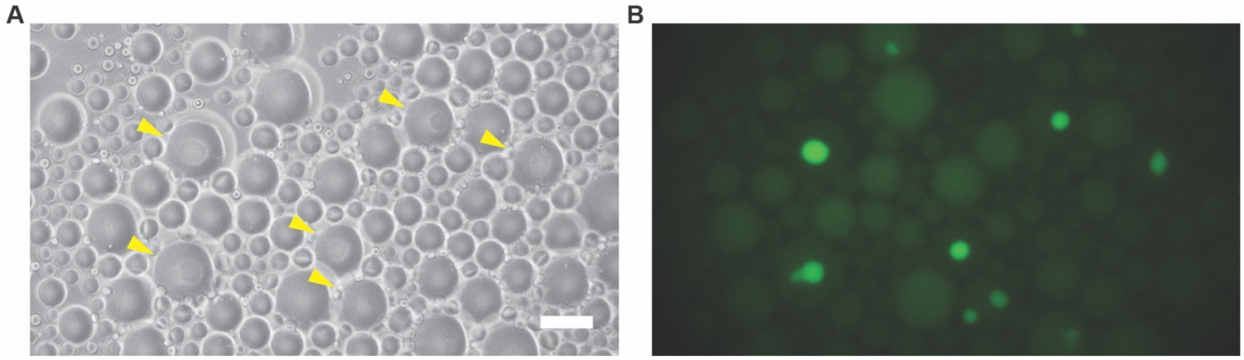


**Fig. S9. Nuclear assembly in water-in-oil emulsions of the egg extract.**

(**A**) Bright field micrograph of water-in-oil emulsions of the egg extract after 90 min incubation at 22ºC. Emulsions with assembled nuclei are highlighted with yellow arrowheads. (**B**) Fluorescence micrograph of the emulsions in GFP channel. GFP-NLS was dispersed in the egg extract prior to emulsion formation for nuclear import assay. Scale bar: 20 μm. See Supplementary Text for the detail of the experiment.

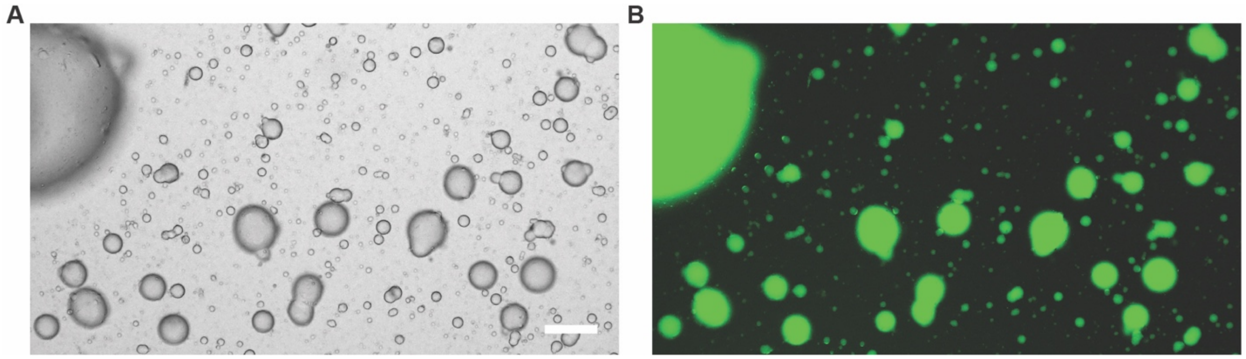


**Fig. S10. Emulsions of the egg extract in lipid-dispersed oil formed by vortex after the incubation of oil-water interface.**

(**A**) Bright field micrograph. (**B**) Fluorescence micrograph of emulsions of the GFP-NLS dispersed egg extract. Condition: $\tau$ = 120 min and $\Delta$ = 10 %. Scale bar: 100 μm.

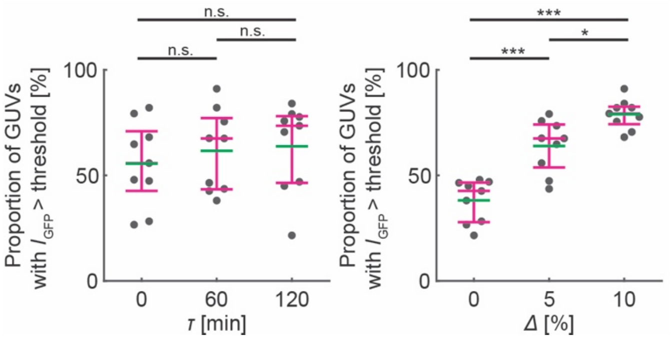


**Fig. S11. Proportion of GUVs with internal mean GFP intensity higher than the background threshold in the observed GUVs (GUVs with aggregated GFP included).**

Results of *post-hoc* Tukey's HSD test after two-way ANOVA focusing on waiting time (left) and chloroform volume (right). Magenta bars indicate Q1-Q3 quartiles and green bars indicate the mean. See "Materials and Methods" in the main text for the details on parameter computation, statistical analysis and Tables S21-23 for the results.

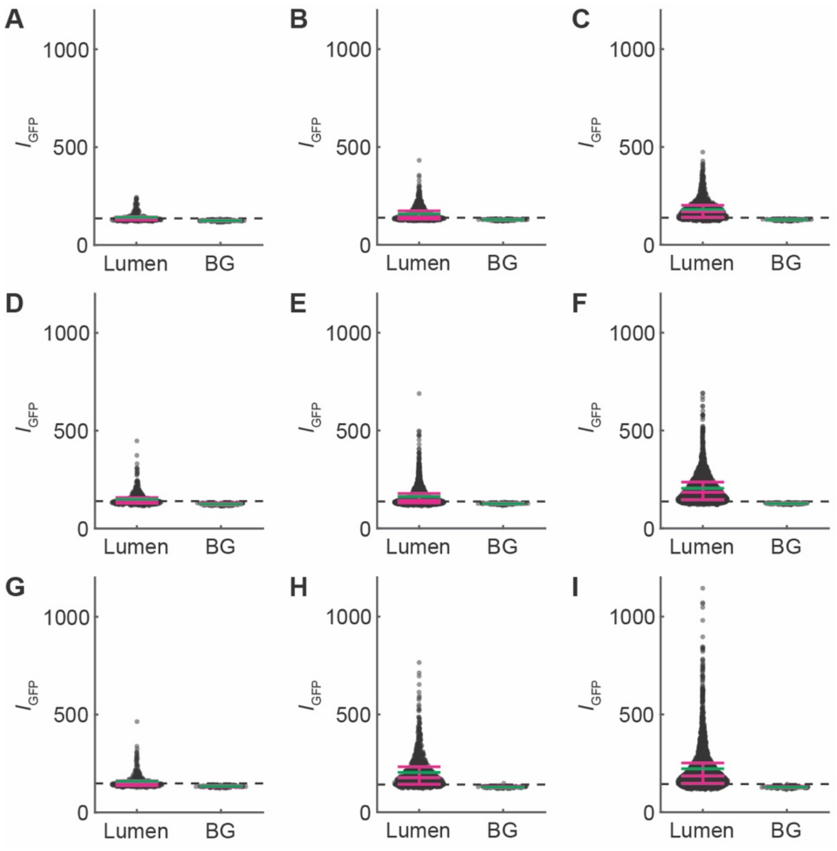


**Fig. S12. Mean GFP intensity in GUVs formed under various conditions of waiting time and chloroform (GUVs with aggregated GFP included).**

(**A**) $\tau = 0$ min, $\Delta = 0$ %. (**B**) $\tau = 0$ min, $\Delta = 5$ %. (**C**) $\tau = 0$ min, $\Delta = 10$ %. (**D**) $\tau = 60$ min, $\Delta = 0$ %. (**E**) $\tau = 60$ min, $\Delta = 5$ %. (**F**) $\tau = 60$ min, $\Delta = 10$ %. (**G**) $\tau = 120$ min, $\Delta = 0$ %. (**H**) $\tau = 120$ min, $\Delta = 5$ %. (**I**) $\tau = 120$ min, $\Delta = 10$ %. "Lumen" denotes the internal mean GFP intensity, $I_{\mathrm{GFP}}$, of GUVs. "BG" denotes the mean GFP intensity of the background at each condition. Magenta bars indicate Q1-Q3 quartiles and green bars indicate the mean. The dashed line indicates the threshold of GFP encapsulation, defined as the mean + 3SD of each background.

| | sum_sq | df | F | PR(>F) |
|---|---|---|---|---|
| C(Waiting_time) | 365706.963 | 2 | 6.945121788 | 0.005814763 |
| C(Chloroform_volume) | 1128924.963 | 2 | 21.43935487 | 1.72689E-05 |
| C(Waiting_time):C(Chloroform_volume) | 257616.5926 | 4 | 2.446191612 | 0.083805462 |
| Residual | 473910 | 18 | | |

**Table S1. Results of the two-way ANOVA for the data in Fig. 2B.**

| | group1 | group2 | meandiff | p-adj | lower | upper | reject |
|---|---|---|---|---|---|---|---|
| 0 | 0 min | 120 min | 233.1111 | 0.1989 | -94.6563 | 560.8785 | FALSE |
| 1 | 0 min | 60 min | 258.6667 | 0.1412 | -69.1008 | 586.4341 | FALSE |
| 2 | 120 min | 60 min | 25.5556 | 0.9793 | -302.2119 | 353.323 | FALSE |

**Table S2. Results of the *post-hoc* Tukey's HSD test for waiting time, as shown in Fig. 2B.**

| | group1 | group2 | meandiff | p-adj | lower | upper | reject |
|---|---|---|---|---|---|---|---|
| 0 | 0% | 10% | 500.7778 | 0.0001 | 249.0645 | 752.4911 | TRUE |
| 1 | 0% | 5% | 242 | 0.0611 | -9.7133 | 493.7133 | FALSE |
| 2 | 10% | 5% | -258.7778 | 0.0431 | -510.4911 | -7.0645 | TRUE |

**Table S3. Results of the *post-hoc* Tukey's HSD test for chloroform volume, as shown in Fig. 2B.**

| Source | Sum Sq | DF | F | PR(>F) |
| --- | --- | --- | --- | --- |
| Waiting_time | 325.2188135 | 2 | 1.848678683 | 0.18611911 |
| Chloroform_volume | 7640.652035 | 2 | 43.4326366 | 1.2935E-07 |
| Waiting_time * Chloroform_volume | 536.7197125 | 4 | 1.525468777 | 0.236959645 |
| Residual | 1583.276395 | 18 | | |

**Table S4. Results of the two-way ANOVA for the data in Figs. 2C.**

| | group1 | group2 | mean_diff | mean_group2 | diff | p-adj | reject |
|---|---|---|---|---|---|---|---|
| 0 | 0 min | 120 min | 55.56410044 | 63.77241689 | -8.208316446 | 0.667984968 | FALSE |
| 1 | 0 min | 60 min | 55.56410044 | 61.58418962 | -6.020089178 | 0.803418371 | FALSE |
| 2 | 120 min | 60 min | 63.77241689 | 61.58418962 | 2.188227268 | 0.971249787 | FALSE |

**Table S5. Results of the *post-hoc* Tukey's HSD test for waiting time, as shown in Figs. 2C.**

| | group1 | group2 | mean_diff | mean_group2 | diff | p-adj | reject |
|---|---|---|---|---|---|---|---|
| 0 | 0% | 10% | 38.15522294 | 78.8965955 | -40.74137255 | 2.74e-08 | TRUE |
| 1 | 0% | 5% | 38.15522294 | 63.8688885 | -25.71366556 | 4.33e-05 | TRUE |
| 2 | 10% | 5% | 78.8965955 | 63.8688885 | 15.02770699 | 0.0114 | TRUE |

**Table S6. Results of the *post-hoc* Tukey's HSD test for chloroform volume, as shown in Figs. 2C.**

| | sum_sq | df | F | PR(>F) |
|---|---|---|---|---|
| C(Waiting_time) | 24422.22222 | 2 | 4.801083411 | 0.021328679 |
| C(Chloroform_volume) | 73813.55556 | 2 | 14.5107613 | 0.000176515 |
| C(Waiting_time):C(Chloroform_volume) | 22931.55556 | 4 | 2.254019105 | 0.103585635 |
| Residual | 45781.33333 | 18 | | |

**Table S7. Results of the two-way ANOVA for the data in Figs. 2E.**

| | group1 | group2 | meandiff | p-adj | lower | upper | reject |
|---|---|---|---|---|---|---|---|
| 0 | 0 min | 120 min | 71.1111 | 0.1447 | -19.6092 | 161.8314 | FALSE |
| 1 | 0 min | 60 min | 52.2222 | 0.3382 | -38.498 | 142.9425 | FALSE |
| 2 | 120 min | 60 min | -18.8889 | 0.8624 | -109.6092 | 71.8314 | FALSE |

**Table S8. Results of the *post-hoc* Tukey's HSD test for waiting time, as shown in Figs. 2E.**

| | group1 | group2 | meandiff | p-adj | lower | upper | reject |
|---|---|---|---|---|---|---|---|
| 0 | 0% | 10% | 120.1111 | 0.0012 | 46.7758 | 193.4465 | TRUE |
| 1 | 0% | 5% | 21.5556 | 0.746 | -51.7798 | 94.8909 | FALSE |
| 2 | 10% | 5% | -98.5556 | 0.0071 | -171.891 | -25.2202 | TRUE |

**Table S9. Results of the *post-hoc* Tukey's HSD test for chloroform volume, as shown in Figs. 2E.**

| | sum_sq | df | F | PR(>F) |
|---|---|---|---|---|
| C(Waiting_time) | 498.1929494 | 2 | 4.307007277 | 0.029611406 |
| C(Chloroform_volume) | 331.6225303 | 2 | 2.866962796 | 0.083009225 |
| C(Waiting_time):C(Chloroform_volume) | 97.8012003 | 4 | 0.422758373 | 0.790171666 |
| Residual | 1041.032962 | 18 | | |

**Table S10. Results of the two-way ANOVA for the data in Figs. 2F.**

| | group1 | group2 | meandiff | p-adj | lower | upper | reject |
|---|---|---|---|---|---|---|---|
| 0 | 0 min | 120 min | 10.4991 | 0.0234 | 1.2844 | 19.7138 | TRUE |
| 1 | 0 min | 60 min | 4.6504 | 0.4306 | -4.5644 | 13.8651 | FALSE |
| 2 | 120 min | 60 min | -5.8487 | 0.2713 | -15.0635 | 3.366 | FALSE |

**Table S11. Results of the *post-hoc* Tukey's HSD test for waiting time, as shown in Figs. 2F.**

| | group1 | group2 | meandiff | p-adj | lower | upper | reject |
|---|---|---|---|---|---|---|---|
| 0 | 0% | 10% | 5.7587 | 0.3184 | -3.964 | 15.4813 | FALSE |
| 1 | 0% | 5% | -2.6342 | 0.7792 | -12.3568 | 7.0885 | FALSE |
| 2 | 10% | 5% | -8.3928 | 0.0998 | -18.1155 | 1.3298 | FALSE |

**Table S12. Results of the *post-hoc* Tukey's HSD test for chloroform volume, as shown in Figs. 2F.**

| | t-statistic | *p*-value | reject |
|---|---|---|---|
| SYTO41 | -0.45551 | 0.64901 | FALSE |

**Table S13. Results of the two-sample t-test for the data in Fig. 4B.**

| | t-statistic | *p*-value | reject |
|---|---|---|---|
| GFP | 21.6719 | 5.37E-68 | TRUE |

**Table S14. Results of the two-sample t-test for the data in Fig. 4C.**

| | sum_sq | df | F | PR(>F) |
|---|---|---|---|---|
| C(Waiting_time) | 1140.75 | 1 | 2.550587 | 0.14892 |
| C(Chloroform_volume) | 3234.083 | 1 | 7.231042 | 0.027537 |
| C(Waiting_time):C(Chloroform_volume) | 990.0833 | 1 | 2.213713 | 0.175107 |
| Residual | 3578 | 8 | | |

**Table S15. Results of the two-way ANOVA for the data in Figs. 4D.**

| | group1 | group2 | meandiff | p-adj | lower | upper | reject |
|---|---|---|---|---|---|---|---|
| 0 | 120 min | 60 min | -19.5 | 0.2544 | -55.4326 | 16.4326 | FALSE |

**Table S16. Results of the *post-hoc* Tukey's HSD test for waiting time, as shown in Figs. 4D.**

| | group1 | group2 | meandiff | p-adj | lower | upper | reject |
|---|---|---|---|---|---|---|---|
| 0 | 10% | 5% | -32.8333 | 0.0386 | -63.5699 | -2.0968 | TRUE |

**Table S17. Results of the *post-hoc* Tukey's HSD test for chloroform volume, as shown in Figs. 4D.**

| Test | H-statistic | *p*-value | reject |
|---|---|---|---|
| Kruskal-Wallis | 87.47423 | 1.01E-19 | TRUE |

**Table S18. Results of the Kruskal-Wallis test for the data in Fig. 4E.**

| | | U-statistic | *p*-value | reject |
|---|---|---|---|---|
| w/ NLC | w/o NLC | 18473.5 | 3.23E-09 | TRUE |
| w/ NLC | w/ nucleus | 563.5 | 6.15E-10 | TRUE |
| w/o NLC | w/ nucleus | 194.5 | 3.41E-14 | TRUE |

**Table S19. Results of the Mann-Whitney U test with Bonferroni correction for the data in Fig. 4E.**

| | | U-statistic | *p*-value | reject |
|---|---|---|---|---|
| NLCs | Nuclei | 1.276622 | 0.201736 | FALSE |

**Table S20. Results of the Wilcoxon rank-sum test (Mann-Whitney U test) for the data in Fig. 4G.**

| Source | Sum Sq | DF | F | PR(>F) |
|---|---|---|---|---|
| Waiting_time | 325.2182186 | 2 | 1.848674936 | 0.186119689 |
| Chloroform_volume | 7640.649423 | 2 | 43.43261317 | 1.29351E-07 |
| Waiting_time * Chloroform_volume | 536.7215111 | 4 | 1.525473587 | 0.23695833 |
| Residual | 1583.276708 | 18 | | |

**Table S21. Results of the two-way ANOVA for the data in Fig. S11.**

| | group1 | group2 | mean_diff | mean_group2 | diff | p-adj | reject |
|---|---|---|---|---|---|---|---|
| 0 | 0 min | 120 min | 55.56411 | 63.77242 | -8.20831 | 0.667985 | FALSE |
| 1 | 0 min | 60 min | 55.56411 | 61.58419 | -6.02008 | 0.803419 | FALSE |
| 2 | 120 min | 60 min | 63.77242 | 61.58419 | 2.188233 | 0.97125 | FALSE |

**Table S22. Results of the *post-hoc* Tukey's HSD test for waiting time, as shown in Fig. S11.**

| | group1 | group2 | mean_diff | mean_group2 | diff | p-adj | reject |
|---|---|---|---|---|---|---|---|
| 0 | 0% | 10% | 38.15523333 | 78.8966 | -40.74136667 | 2.74e-08 | TRUE |
| 1 | 0% | 5% | 38.15523333 | 63.86888889 | -25.71365556 | 4.33e-05 | TRUE |
| 2 | 10% | 5% | 78.8966 | 63.86888889 | 15.02771111 | 0.0114 | TRUE |

**Table S23. Results of the *post-hoc* Tukey's HSD test for chloroform volume, as shown in Fig. S11.**
